# RUNX1::RUNX1T1 Depletion Eliminates Stemness and Induces Bidirectional Differentiation of Acute Myeloid Leukemia

**DOI:** 10.1101/2025.02.13.637937

**Authors:** Polina K Derevyanko, Laura E. Swart, L. Daniel Mata Casimiro, Luca van den Brink, Anita T. van Oort, Minoo Ashtiani, C. Michel Zwaan, Anja Krippner-Heidenreich, Constanze Bonifer, Raymond Schiffelers, H. Josef Vormoor, Sophie G. Kellaway, Olaf Heidenreich

**Affiliations:** Princess Máxima Center for Pediatric Oncology, Utrecht, The Netherlands; Department of Pediatric Hematology/Oncology, Erasmus MC Sophia Children’s Hospital, Rotterdam, The Netherlands; Institute of Cancer and Genomic Sciences, University of Birmingham, Birmingham, UK; Murdoch Children’s Research Institute, Royal Children’s Hospital, Australia; Department of Pharmaceutics, Utrecht Institute for Pharmaceutical Sciences, Utrecht University, The Netherlands; Blood Cancer and Stem Cells, Centre for Cancer Sciences, School of Medicine, University of Nottingham, Nottingham, UK; Department of Hematology, University Medical Center Utrecht, The Netherlands; Wolfson Childhood Cancer Research Centre, Translational and Clinical Research Institute, Newcastle University, Newcastle upon Tyne, UK

**Author notes:** Authors contributed equally (Polina K Derevyanko, Laura E. Swart).

**Keywords:** Acute myeloid leukemia, RUNX1::RUNX1T1, siRNA delivery, primary AML cells, leukemic differentiation

## Abstract

Chromosomal rearrangements that generate novel fusion genes are a hallmark of acute myeloid leukemia (AML). Depletion experiments in cell line models have suggested that continued expression is required for maintaining their leukemic phenotype and they therefore represent ideal cancer-specific therapeutic targets. However, to which extent this result holds true for the different stages of hematopoietic development in primary cells and whether therapeutic agents can be efficiently delivered to those cells is still unclear. In this study, we demonstrate that primary AML cells harboring the chromosomal translocation t(8;21) are critically dependent on the corresponding fusion gene, *RUNX1::RUNX1T1*, to suppress differentiation and maintain stemness. Silencing *RUNX1::RUNX1T1* expression using siRNA-loaded lipid nanoparticles induces substantial changes in chromatin accessibility, thereby redirecting the leukemia-associated transcriptional network towards a myeloid differentiation program. Single-cell analyses reveal that this transcriptional reprogramming is associated with the depletion of immature stem and progenitor-like cell populations, accompanied by an expansion of granulocytic and eosinophilic/mast cell-like populations with impaired self-renewal capacity. These findings underscore the essential role of *RUNX1::RUNX1T1* in sustaining AML and highlight the therapeutic potential of targeting fusion gene expression in primary AML cells.

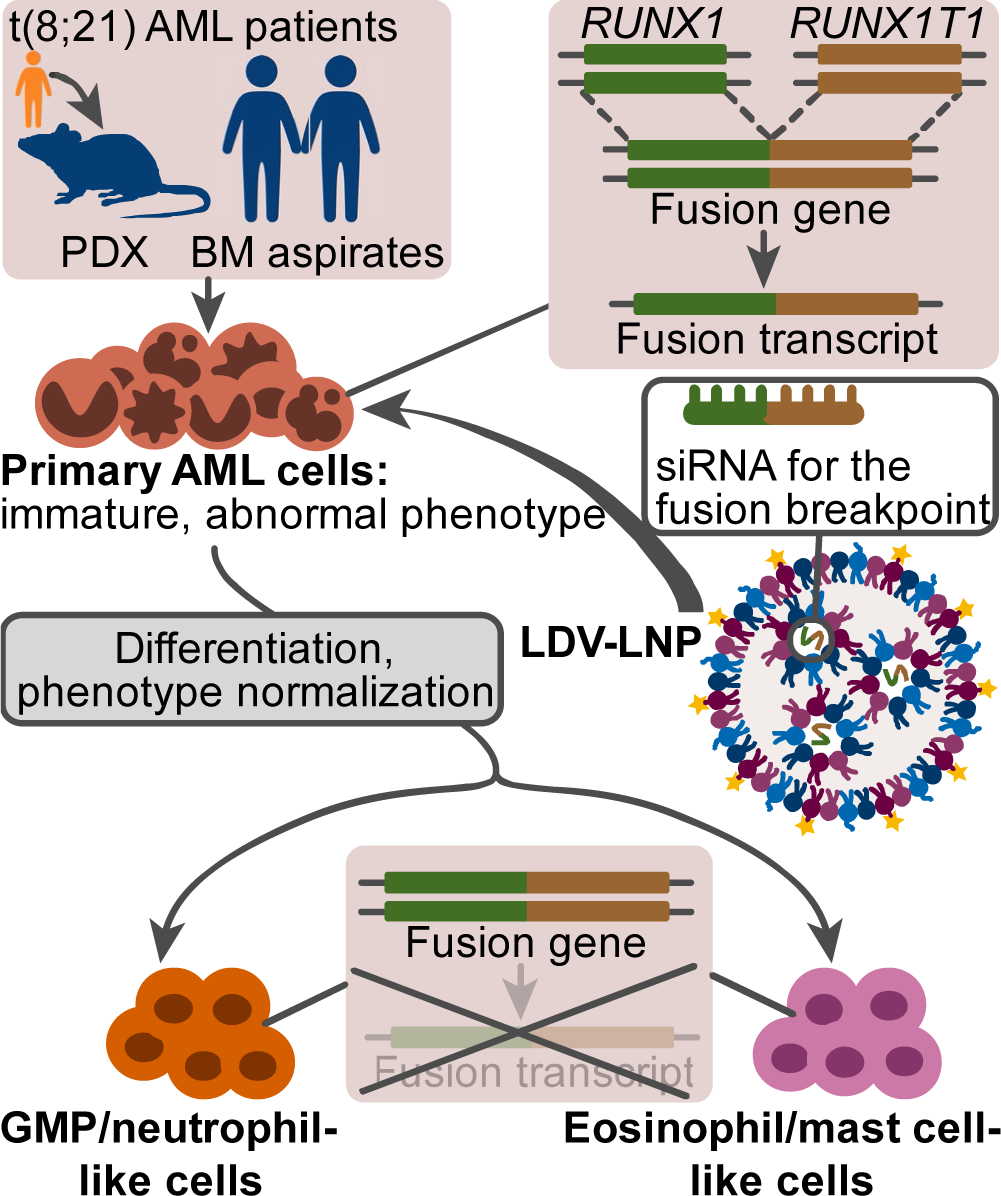

## Introduction

Current treatment for acute myeloid leukemia (AML) relies heavily on intensive chemotherapy, which is associated with significant toxicities, profoundly affecting patients’ quality of life ^1,2^ ^3^. Despite these aggressive regimens, a significant proportion of 60–70% of adult AML patients aged 60 years and younger ultimately succumb to the disease ^4^. Pediatric patients have a better prognosis, with average relapse rates of 25% ^5^. However, due to their longer life expectancy, they are disproportionately affected by the long-term adverse effects of chemotherapy ^6,7^. This highlights the urgent need for more targeted, leukemia-specific therapies for AML.

A hallmark of AML are various chromosomal rearrangements, which result in the expression of leukemia-specific fusion proteins in approximately 30% of adult AML cases and over 50% of pediatric AML cases ^8,9^. Expression of these proteins is leukemia-initiating and is required to maintain the leukemogenicity of AML cells, thus representing highly attractive therapeutic targets ^10^. However, many AML fusion genes encode transcriptional regulators, which are notoriously challenging to target using conventional drug design due to a lack of stably structured, druggable regions. Moreover, these regions are often shared with the wild-type protein encoded by the intact allele, leading to undesirable on-target toxicities ^11^.

Among AML fusion genes, RUNX1::RUNX1T1 is the most prevalent recurrent genetic aberration in pediatric and younger adult AML, occurring in 10–15% of cases ^12^. Prior studies have highlighted the role of RUNX1::RUNX1T1 in driving leukemic self-renewal and impeding differentiation in cell lines and transgenic models ^13–16^. However, these models fail to fully capture the intra-tumoral and inter-patient heterogeneity of AML. The role of leukemic fusion genes in primary patient-derived AML cells remains poorly understood, largely due to the experimental challenges in perturbing fusion gene expression specifically within these cells.

Critical questions remain unanswered: How does the loss of fusion gene expression affect the viability, differentiation, and self-renewal of various AML cell populations, and how do these effects vary between patients? What differentiation pathways are activated upon gene silencing, and can restoration of fusion gene expression reverse differentiation? Is there a "point of no return" where AML cells lose stemness entirely, or can they retain or regain self-renewal despite adopting a differentiated phenotype?

Addressing these questions necessitates tools that can specifically inhibit fusion gene expression in primary AML cells. Previous work, including ours, has demonstrated the feasibility of targeting fusion genes using siRNAs designed to specifically bind the fusion junction site ^13,17–19^. Encapsulation of these siRNAs in lipid nanoparticles facilitates efficient and specific knockdown of fusion gene expression in cell lines and patient-derived cells, both in vitro and in xenotransplanted mice ^20–22^. Leveraging the power of this system, we have now applied ex vivo manipulation of patient-derived AML cells to study the impact of fusion gene knockdown. Our data reveal a strict dependency of RUNX1::RUNX1T1-expressing AML cells on continuous fusion gene expression. Even temporary silencing of the gene exerts profound and lasting effects on leukemic self-renewal and differentiation, underscoring the therapeutic potential of targeting fusion genes in AML.

## Results

### A lipid nanoparticle formulation for gene knockdown in patient-derived AML cells

Primary AML cells are notoriously hard to transfect. To perturb gene expression in both primary and patient-derived xenograft (PDX) AML cells, we therefore developed targeted lipid nanoparticles (LNPs) designed for the delivery of *RUNX1::RUNX1T1* and mismatch control siRNAs (Fig. 1A, B) to specific cells. In previous work, we demonstrated that siRNA-loaded LNPs effectively suppressed *RUNX1::RUNX1T1* expression in t(8;21)-positive AML cell lines, both in vitro and in xenotransplanted immunodeficient mice ^21^. To enhance siRNA delivery and efficacy in primary AML cells, we functionalized LNPs with a high-affinity ligand, 4-((N’-2-methylphenyl)ureido)-phenylacetyl-L-leucyl-L-α-aspartyl-L-valyl-conjugate (LDV), targeting the Very Late Antigen-4 (VLA-4), an integrin α4-β1 heterodimer that is commonly expressed on hematopoietic cells but rarely on other lineages ^24,25,57^.

**Figure 1.**
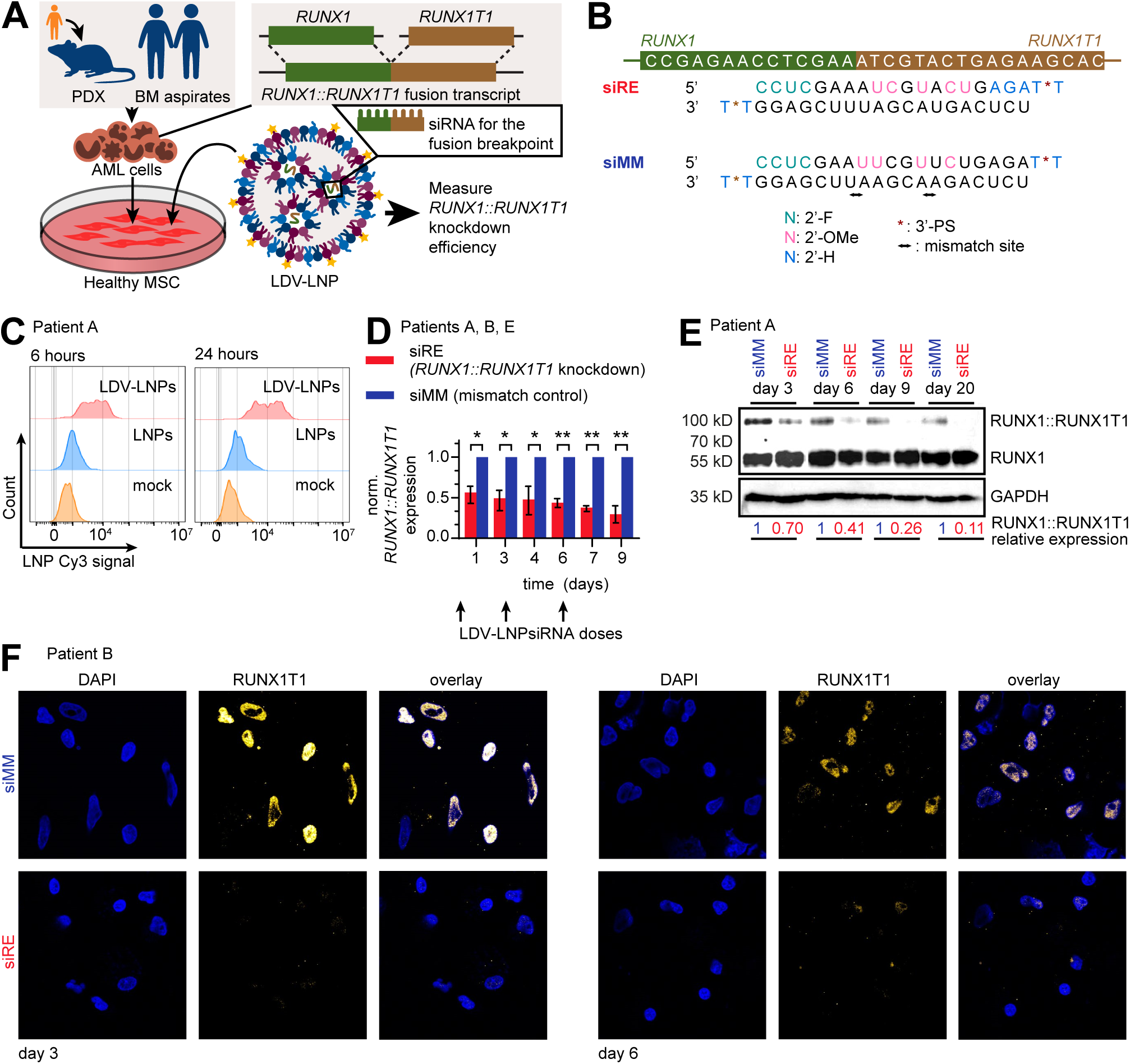
LDV-LNPs have improved uptake and siRNA delivery efficacy in t(8;21) positive primary AML cells. (A) Schematic illustration of the experimental approach. (B) Chemically modified siRNAs (siRE, siMM) were made by introducing of 2’-fluoro- (2’-F), 2’-methoxy (2’-OMe) and 2’-deoxy- (2’-H) and 3’-terminal phosphorothioate (PS). siRE is complimentary to the RUNX1::RUNX1T1 fusion breakpoint; siMM contains 2 mismatch sites. (C) Histogram showing LNP Cy3 signal in a RUNX1::RUNX1T1-positive primary sample after 6 hours (left) or 24 hours (right) for LDV-LNPs (red, top), LNPs (blue, middle) and PBS (orange, bottom) as measured by flow cytometry. (D, E) Reduction of the *RUNX1::RUNX1T1* fusion transcript (D) and protein (E) of siRNA LNP-treated patient cells detected by qPCR or western blotting after sequential LNP dosing. (D) Mean + SD are displayed. *P <0.05, **P < 0.005. n = 3 different t(8;21) primary patient bone marrow aspirates, one-tailed one-sample t-test. (F) Images showing intracellular RUNX1T1 detection by confocal microscopy after 3 (left) and 6 days (right) in primary patient B cells incubated with siMM-mod LDV-LNPs (top panel) or siRE-mod LDV-LNPs (bottom panel). siRE, *RUNX1::RUNX1T1* siRNA; siMM, mismatch control.

We evaluated LNP uptake and siRNA efficacy in primary AML cells co-cultured with human bone marrow mesenchymal stromal cells (MSCs) under serum-free conditions (Fig. 1A). LDV-functionalized LNPs (LDV-LNPs) exhibited a 5-fold and 9-fold improvement in uptake in primary AML cells at 6 and 24 hours, respectively, compared to non-functionalized LNPs (Fig. 1C, Suppl. Fig. 1A, B). Uptake was neither patient-specific nor fusion gene-dependent, as demonstrated by the improved uptake observed in additional normal karyotype AML bone marrow aspirate sample (Suppl. Fig. 1B).

Using the RUNX1::RUNX1T1 fusion transcript as the target, we encapsulated an siRNA complementary to the fusion site of *RUNX1::RUNX1T1* mRNA and monitored its knockdown at both transcript and protein levels in primary AML cells (Fig. 1A, B). Treatment of three *RUNX1::RUNX1T1*-positive AML patient samples and one PDX sample with LDV-LNPs, administered on days 0, 3, and 6 of co-culture, resulted in a sustained 2- to 3-fold reduction in *RUNX1::RUNX1T1* transcripts and a greater than 3-fold reduction in the corresponding fusion protein levels (Fig. 1D, E, Suppl. Fig. 1C, D). Depletion of the RUNX1::RUNX1T1 protein persisted for over 10 days following the final LNP treatment (Fig. 1E). In contrast, expression of wild-type RUNX1 protein, which is not expressed from the *RUNX1::RUNX1T1* fusion allele, remained unaffected, underscoring the precision of the siRNA in targeting the fusion transcript (Fig. 1E). To confirm the specificity of the knockdown in primary cells, we performed intracellular staining for RUNX1T1, revealing a marked loss of the nuclear fusion protein (Fig. 1F).

In summary, these findings demonstrate the feasibility of effectively perturbing fusion gene expression *ex vivo* in primary AML cells.

### Silencing of RUNX1::RUNX1T1 alters chromatin accessibility

To investigate the impact of RUNX1::RUNX1T1 modulation on chromatin accessibility, we performed ATAC-seq on AML PDX cells, identifying 57,339 distal open chromatin sites. *RUNX1::RUNX1T1* knockdown altered chromatin accessibility in 6.9% (3,953) and 4.5% (2,596) of these sites, increasing and decreasing accessibility, respectively (Fig. 2A, Suppl. Fig. 2A–D). Sites with increased accessibility were enriched for C/EBP motifs, while those with decreased accessibility were enriched for AP-1 motifs (Fig. 2B, C). Both AP-1 and C/EBP family transcription factors are pivotal in myeloid differentiation ^58–60^. Moreover, AP-1 motifs are associated with genes relevant for leukemic proliferation in t(8;21) AML ^14,23,47^. Comparing these chromatin changes to lineage-specific profiles revealed that regions losing accessibility corresponded to HSC/MPP-specific patterns, while newly accessible regions reflected GMP, monocyte, and neutrophil-specific profiles (Fig. 2D).

**Figure 2.**
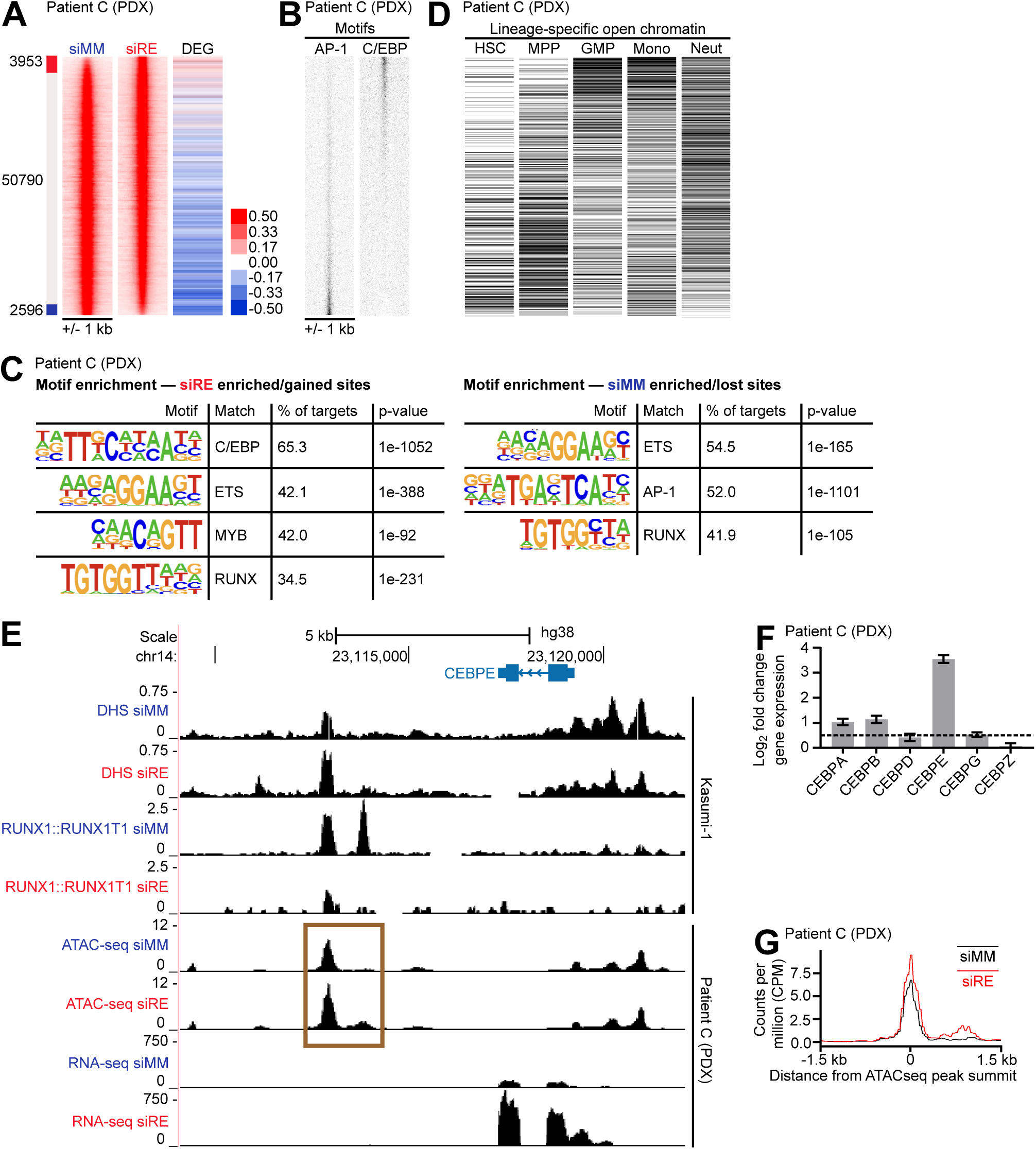
RUNX1::RUNX1T1 depletion leads to global chromatin changes in a t(8;21) AML PDX. (A) Heatmaps depicting the chromatin accessibility in siMM and siRE-treated PDX samples. Distal (>1.5kb from TSS) ATAC-seq sites were ranked by the fold change between siRE and siMM. N=3 replicates. The bar on the left indicates the significantly (adj.p<0.05, fold change >2) gained (red bar) and lost (blue bar) peaks. (B) AP-1 and C/EBP binding motifs plotted across the same scale as (A). (C) Lineage-specific open chromatin sites plotted across the same scale as (A) as a binary, black and white indicate open chromatin sites specific or not specific for that lineage. (C) De novo motif discovery on significantly lost/gained sites in siMM and siRE-treated PDX samples. (D) Lineage-specific open chromatin sites plotted across the same scale as (A) as a binary, black and white indicate open chromatin sites specific or not specific for that lineage. (E) Bar graph showing the log2fold change gene expression for the C/EBP family genes. (F) UCSC Genome Browser screenshot showing *CEBPE* locus. Upper four tracks show DNase1-seq and RUNX1::RUNX1T1 ChIP-seq ^48^, lower four tracks show patient C PDX ATAC-seq and RNA-seq. (G) Average profile of ATAC-seq read depth (counts per million) of merged replicates at the differentially accessible peak indicated in (F). siRE, *RUNX1::RUNX1T1* siRNA; siMM, mismatch control.

The C/EBP transcription factor family shares a common DNA-binding motif and is critical for granulocytic differentiation ^61–63^. Repression of C/EBPα is associated with the differentiation block in t(8;21) AML ^64^, and *CEBPE* is a direct target of RUNX1::RUNX1T1-mediated repression (Fig. 2E) ^48,65^. *RUNX1::RUNX1T1* knockdown upregulated *CEBPA*, *CEBPB*, and *CEBPE* transcript levels and increased accessibility of C/EBPε-associated cis-regulatory regions (Suppl. Table 6, Fig. 2E–G). Integration with ChIP-seq and DHS-seq data from the t(8;21) AML cell line Kasumi-1 confirmed that RUNX1::RUNX1T1 depletion alters the same chromatin sites in a similar manner as in patient-derived cells, directly affecting RUNX1::RUNX1T1-occupied regions and leading to increased *CEBPA* and *CEBPE* expression and occupation of differentiation associated loci such as *CTSG* and *RNASE2* and the immune checkpoint gene *VSIR* (Suppl. Fig. 2E–G) ^15,66^.

In summary, these data demonstrate that RUNX1::RUNX1T1 depletion (i) abolishes a stem-cell-specific chromatin pattern and (ii) alleviates the differentiation block in t(8;21) AML by activating a C/EBPα- and C/EBPε-driven transcriptional program, thereby promoting myeloid differentiation.

### RUNX1::RUNX1T1 depletion activates differentiation-linked transcriptional programs

To correlate gene expression changes with chromatin status in patient-derived AML cells, we conducted bulk RNA sequencing (RNA-seq) on t(8;21) AML PDX cells ^23^ following *RUNX1::RUNX1T1* knockdown. Principal component analysis (PCA) of bulk RNA-seq results robustly separated LDV-LNPsiRE-treated cells from LDV-LNPsiMM-treated controls (Suppl. Fig. 3A). Previous studies in AML cell lines have demonstrated that RUNX1::RUNX1T1 knockdown shifts transcriptional programs from self-renewal to differentiation, while its expression in hematopoietic stem and progenitor cells (HSPCs) promotes self-renewal ^14,15,21,47,48,67^. Gene set enrichment analysis (GSEA) confirmed this pattern in RUNX1::RUNX1T1-depleted AML PDX cells (Fig. 3A, Suppl. Fig. 3B, C). Differential gene expression analysis identified 576 upregulated and 941 downregulated genes (adjusted p-value < 0.001; log2-fold change > 1; base mean ≥ 50) (Fig. 3B, C, Suppl. Fig. 3D, Suppl. Table 6). Among these were known transcriptional targets of RUNX1::RUNX1T1 linked to granulocytic and neutrophilic differentiation (e.g., *FUT7, ELANE* and *CTSG*) and *VSIR* encoding an immune checkpoint factor (Fig. 3B, C) ^68–70^.

**Figure 3.**
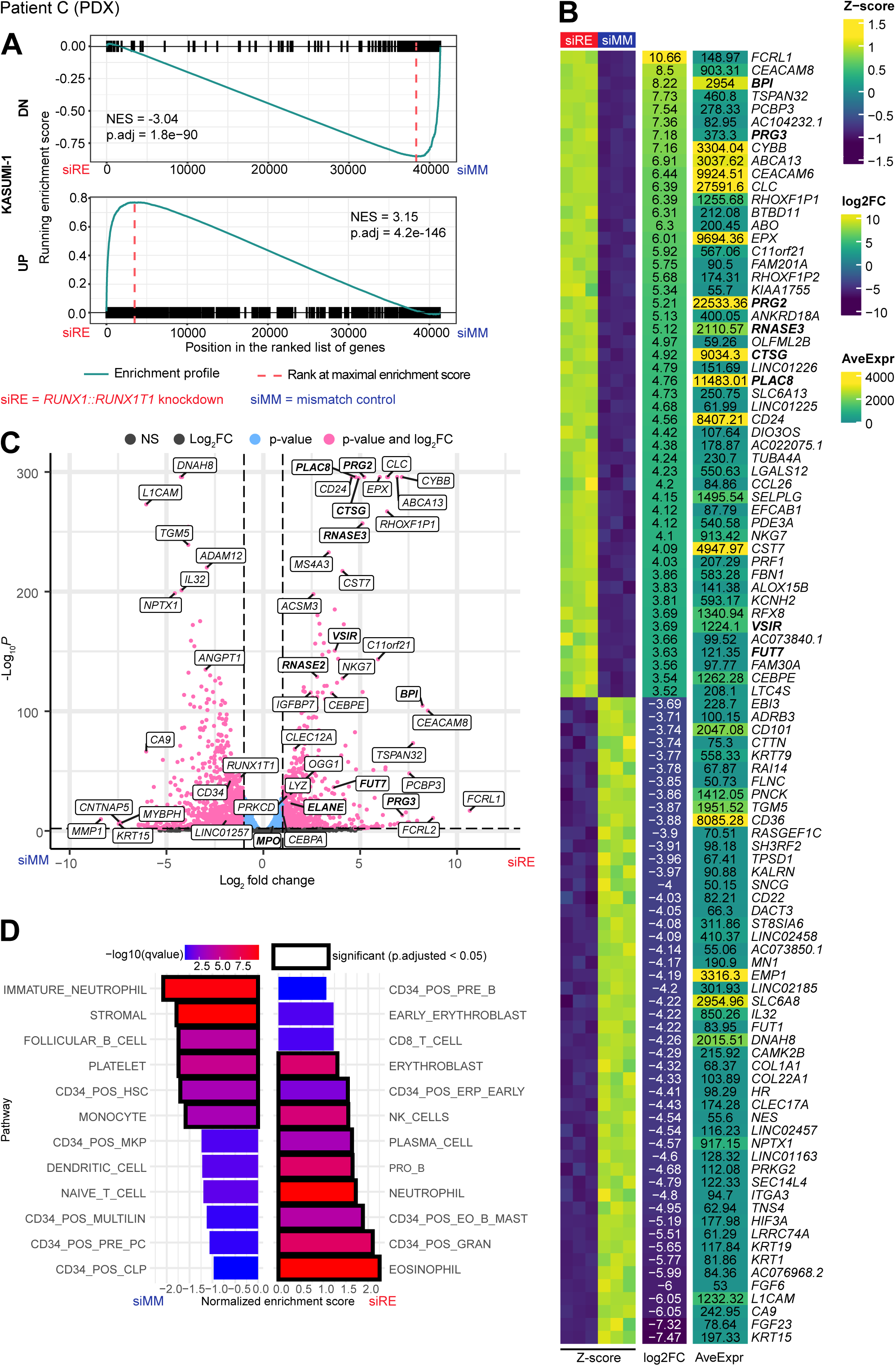
RUNX1::RUNX1T1 depletion in PDX AML cells leads to global transcriptome changes. (A) GSEA plots for the bulk RNA-seq of siRNA-treated PDX samples against gene sets of differentially (log2FC < −1 or log2FC > 1, adj. p < 0.05) down- (top panel) or upregulated (bottom panel) genes in siRNA-treated Kasumi-1 cells (data from ^21^). (B) Heatmap showing the Z-scores, log2FCs and normalized counts (DESeq2 normalization) for 50 genes with the highest and lowest log2FC between siRE and siMM-treated PDX samples. (C) Volcano plot of differential expression analysis (DESeq2) of siRNA-treated PDX samples. Adjusted p-value cutoff = 0.001. Log2FC cutoff = 1. (D) Normalized enrichment scores for pathways from the Human Cell Atlas bone marrow gene set collection ^55^ of siRNA-treated PDX samples. Positive enrichment scores correspond to an enrichment in the samples treated with siRE LDV-LNPs. siRE, *RUNX1::RUNX1T1* siRNA; siMM, mismatch control.

Intriguingly, *RUNX1::RUNX1T1* silencing also upregulated eosinophil-associated genes (*RNASE2, RNASE3, PRG2, PRG3*; Fig. 3B, C), suggesting differentiation potential toward neutrophilic and eosinophilic lineages. GSEA for bone marrow cell type markers supported this observation, showing a shift from a transcriptome characteristic for immature cells to those of granulocytes, eosinophils, and mast cells (Fig. 3D). Collectively, these results demonstrate that the loss of RUNX1::RUNX1T1 activates transcriptional programs promoting granulocytic and eosinophilic differentiation.

### Loss of RUNX1::RUNX1T1 drives differentiation of primary AML cells

To investigate the effects of LNP treatment and RUNX1::RUNX1T1 depletion on primary cell subpopulations and their fate in patient-derived AML, we performed single-cell RNA sequencing (scRNA-seq) on PDX and two RUNX1::RUNX1T1-positive primary bone marrow aspirates treated with siRE or siMM-carrying LNPs (Suppl. Fig. 4A, Suppl. Table 7). Following quality control, filtering, and exclusion of MSCs through SNP-based demultiplexing and MSC marker gene analysis (Suppl. Fig. 4B–D), we obtained 8,732, 7,021, and 3,108 cells from AML patients A, B, and C (PDX), respectively. *RUNX1T1* expression is substantially lower in normal HSPCs compared to the fusion gene and was therefore used as a surrogate marker for RUNX1::RUNX1T1 to identify leukemic cells ^15^. Four smaller clusters in patient A and two in patient B lacked fusion gene expression under both conditions, representing normal hematopoietic cells. As expected, no human non-leukemic clusters were present in the patient C PDX sample (Fig. 4B, C).

**Figure 4.**
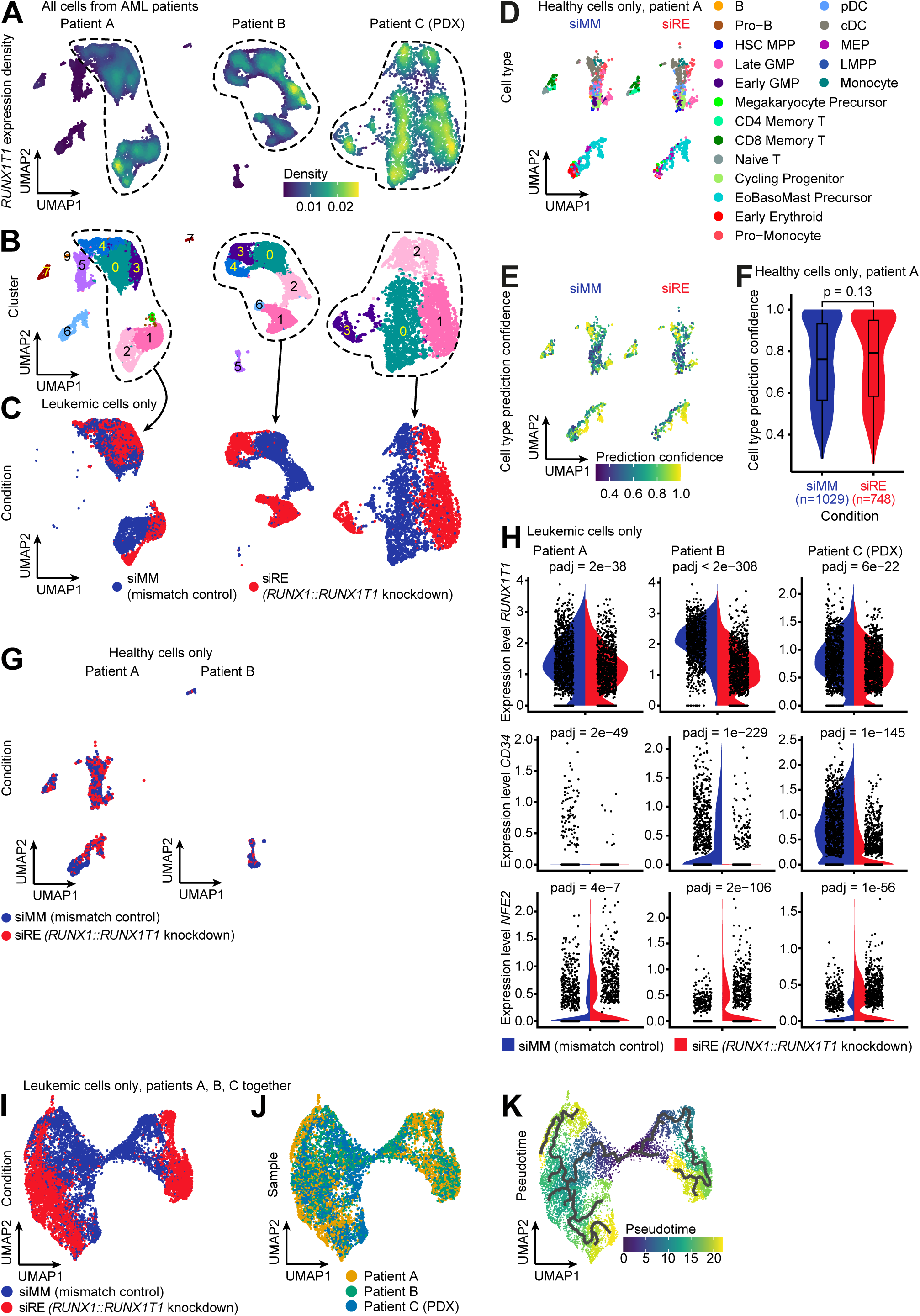
*RUNX1::RUNX1T1* silencing drives differentiation of primary AML cells without affecting normal cell populations. (A) UMAP plots showing *RUNX1T1* expression density in individual patients. (B) UMAP plots of the AML patient cells, colored by cluster. (C) UMAP plot showing distribution of siRE (red) and siMM (blue) treated AML cells. (D) UMAP plots of normal bone marrow cells from patient A, colored by predicted cell type. (E) UMAP plots of normal bone marrow cells from AML patient A, colored by cell type prediction confidence. (F) Box and violin plot of confidence scores for annotation of normal bone marrow cells from AML patient A. Wilcoxon rank sum test was used to determine the p-value. (G) UMAP plot showing distribution of siRE (red) and siMM (blue) treated normal bone marrow cells from two patients. (H) Violin plots of *RUNX1::RUNX1T1* (top panel) and its direct target genes *CD34* (middle panel) and *NFE2* (bottom panel) in scRNAseq data of the 2 patient samples and 1 AML PDX. Wilcoxon rank sum test with Bonferroni correction was used for p-value determination. (I, J) UMAP plot of integrated scRNAseq dataset of 3 patients colored by experimental condition (I) or patient (J). (K) UMAP plot showing pseudotime graph, rooted at the cell population with the highest HSC module score (as depicted in Fig. 7B). siRE, *RUNX1::RUNX1T1* siRNA; siMM, mismatch control.

Analysis of non-leukemic clusters shown in the UMAP from both LDV-LNP-siRE-treated and LDV-LNP-siMM-treated conditions revealed the presence of HSPCs, myeloid and erythroid progenitors, B, T, and NK cells (Fig. 4D). LNP-mediated siRE delivery did not affect normal hematopoietic cell fates as transcriptomic analysis showed no significant differences between the two conditions (Fig. 4E–G, Suppl. Fig. 4E, G). siRE-LNP treatment induced robust *RUNX1::RUNX1T1* knockdown in leukemic cells, demonstrated by reduced *RUNX1T1* reads (Fig. 4H, top row). Here, knockdown led to significant transcriptional changes in RUNX1::RUNX1T1 target genes, including the downregulation of the HSPC marker *CD34* and upregulation of *NFE2*, a key driver of erythroid, megakaryocytic and granulocytic differentiation (Fig. 4H, middle and bottom rows). The transcriptomic shift following *RUNX1::RUNX1T1* silencing (Fig. 4C) was consistent with shifts observed in bulk RNAseq of AML PDX samples (Suppl. Fig. 4F).

Integration of altered gene expression across the three patient samples identified 216 significantly upregulated and 7,403 downregulated genes (adjusted p < 0.001) following *RUNX1::RUNX1T1* silencing, indicating a substantial reorganization of transcriptional networks (Suppl. Table 7). This reorganization was reflected in the UMAP and PCA, which revealed a pronounced shift in the composition of the integrated cell populations after knockdown (Fig. 4I, J, Suppl. Fig. 4H). Interestingly, pseudotime analysis suggested this shift exhibited different directions and that this bi-directional myeloid differentiation originates at the intersection of the two most immature cell clusters, which are likely to harbor the LSCs (Fig. 4K). In conclusion, loss of RUNX1::RUNX1T1 facilitates differentiation of immature leukemic cells along distinct cell fates.

### RUNX1::RUNX1T1 depletion enhances bidirectional granulocytic and eosinophilic differentiation

To characterize the differentiation directions, we examined the gene expression patterns in the cells belonging to the two groups. Cluster analysis identified seven distinct clusters, with two central clusters (clusters 2 and 4) disappearing upon *RUNX1::RUNX1T1* depletion (Fig. 5A). The remaining five clusters were shared between control and knockdown cells. However, *RUNX1::RUNX1T1* knockdown caused an additional shift along UMAP, away from the center. These data indicate that the AML cells do undergo a limited differentiation even in the presence of the fusion and that silencing of *RUNX1::RUNX1T1* causes a further progression along two different lineages.

**Figure 5.**
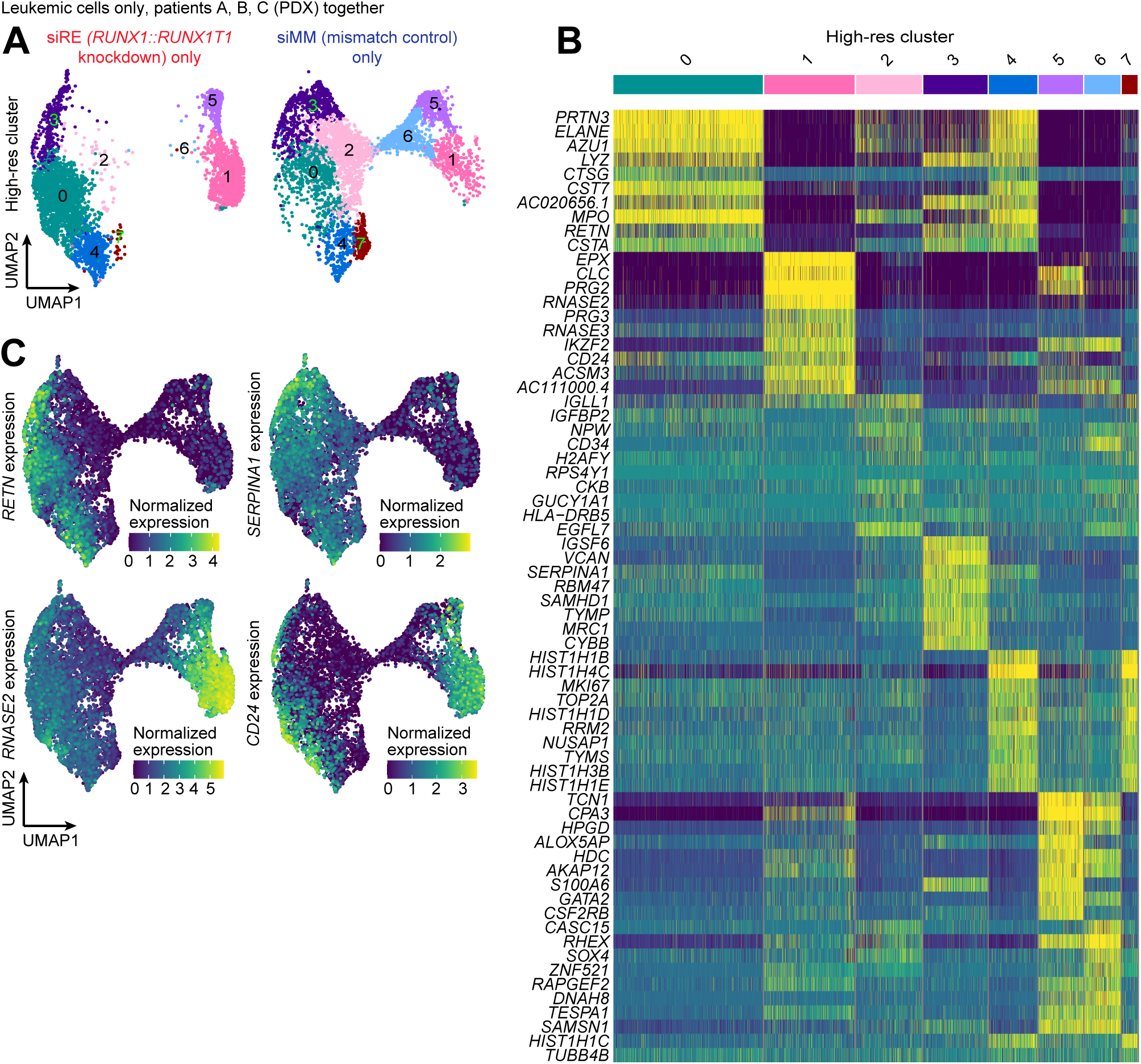
Depletion of RUNX1::RUNX1T1 enhances bidirectional granulocytic and eosinophilic differentiation. (A) UMAP plots of the integrated scRNAseq dataset, split by experimental condition and colored by cluster. (B) Heatmap of top marker genes for each cluster depicted in (A). (C) UMAP plots of the integrated scRNAseq dataset showing expression distribution of myeloid marker genes. siRE, *RUNX1::RUNX1T1* siRNA; siMM, mismatch control.

These shifts were associated with increased expression of granulocytic genes (e.g., *ELANE, CTSG, RETN*, and *SERPINA1*) in clusters 0, 3, and 4, and eosinophilic genes (e.g., *PRG2, CLC,* and *RNASE2*) in clusters 1, 5, and 6 (Fig. 5B, C). In contrast, expression of *CD24*, a marker associated with the transition from the promyelocytic to the myelocytic stage and a “don’t eat me” signal for macrophages, increased substantially in all cell populations with *RUNX1::RUNX1T1* knockdown (Fig. 5C) ^71,72^, highlighting advanced granulocytic and eosinophilic differentiation. These findings demonstrate that RUNX1::RUNX1T1 knockdown leads to enhanced differentiation toward granulocytic and eosinophilic lineages, and the loss of immature leukemic populations.

To examine the differentiation patterns observed in *RUNX1::RUNX1T1*-silenced leukemic cell populations in more detail, we projected our scRNA-seq datasets onto a human hematopoiesis reference map comprising 55 distinct cell states ^54^. This analysis identified 11 leukemic states resembling normal hematopoietic cell states (Fig. 6A). Leukemic cells with transcriptional profiles resembling megakaryocytic/erythrocytic progenitors (MEPs), early granulocyte-macrophage progenitors (GMPs), and lymphoid progenitors were located in the joint region of the UMAP, which was lost upon *RUNX1::RUNX1T1* silencing. These immature cell states were bordered by late GMP, pro-monocytic, and monocytic states on one side, and eosinophilic/basophilic/mast cell precursors on the other (Fig. 6A). Projection confidence scores were significantly higher for *RUNX1::RUNX1T1*-silenced cells compared to control cells for both directions of differentiation (Fig. 6B, Suppl. Fig. 5A) suggesting a shift from an aberrant limited differentiation to a more normal differentiation upon RUNX1::RUNX1T1 loss.

**Figure 6.**
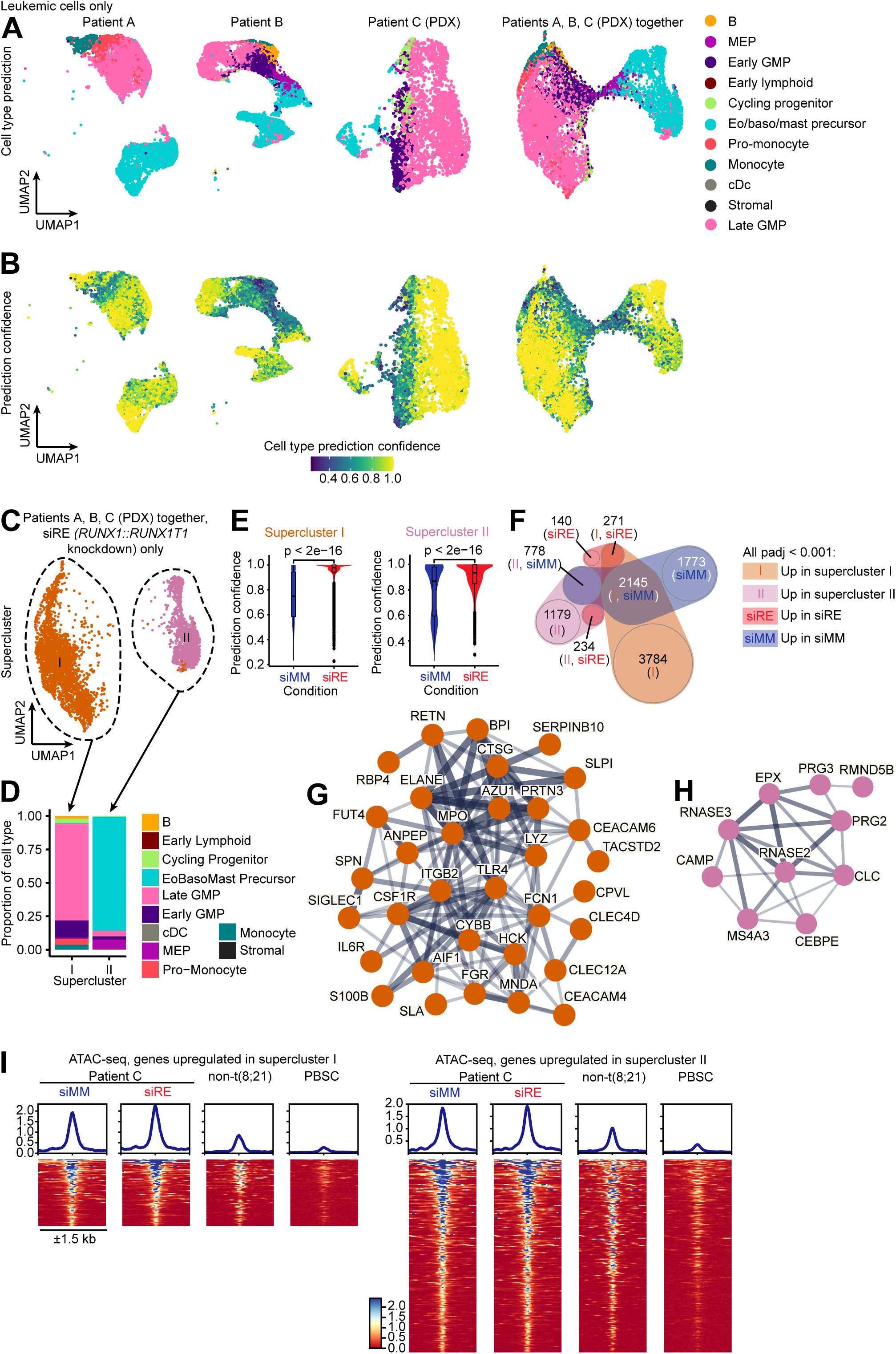
Depletion of RUNX1::RUNX1T1 enhances bidirectional granulocytic and eosinophilic differentiation. (A– B) UMAP plots of scRNAseq data of the 3 patients separately (left, middle left, middle right) and integrated (right), colored by predicted cell type (A) or cell type prediction confidence score (B). (C) UMAP plot of integrated scRNAseq data of the 3 patients, siRE condition only. (D) Proportions of predicted cell types in the 2 superclusters depicted in (C). (E) Box and violin plots showing cell type prediction confidence score in scRNAseq data of the 3 patients with and without RUNX1::RUNX1T1 depletion for supercluster I (granulocytic differentiation, left) and supercluster II (eosinophilic differentiation, right). (F) Venn diagram depicting significantly (adj. p < 0.001) upregulated genes in integrated scRNAseq data of the 3 patients, supercluster 1 (orange), supercluster II (pink), siRE condition (red), siMM condition (blue). (G, H) STRING-generated gene networks of clusters formed by genes upregulated upon *RUNX1::RUNX1T1* silencing and higher expressed in supercluster I (G) or supercluster II (H). (I) Heatmaps with associated average profiles showing normalised read counts (counts per million) of patient C (PDX) merged replicates ATAC-seq and DNaseI-seq from a non-t(8;21) AML patient and healthy CD34+ sample ^47^. Left panel shows coverage at accessible chromatin sites found in patient C (PDX) associated with supercluster I (n=73 sites) and the right panel at sites associated with supercluster II (n=212) marker genes (RUNX1::RUNX1T1 knockdown only, >0.5-fold difference, adjusted p-value <0.05). siRE, *RUNX1::RUNX1T1* siRNA; siMM, mismatch control.

To further investigate how fusion gene knockdown drives bidirectional differentiation, we grouped the *RUNX1::RUNX1T1*-depleted cells into two superclusters (Fig. 6C). Supercluster I, encompassing parts of clusters 0, 3, and 7, predominantly exhibited a late GMP-like phenotype adopted by ∼75% of the cells (Figs. 5A, 6D). Conversely, nearly 90% of cells in supercluster II, comprising parts of clusters 1 and 5, displayed transcriptional profiles resembling eosinophilic/basophilic/mast cell precursors (Fig. 6D). Cell type projection confidence scores were significantly higher for AML cells upon RUNX1::RUNX1T1 depletion in both superclusters (Fig. 6E), indicating that the shift from an aberrant limited differentiation to a more normal differentiation happened along both differentiation trajectories. These two differentiation trajectories were further corroborated by the analysis of cell type module scores between the two superclusters (Suppl. Fig. 5B).

Next, we asked which genes that are differentially expressed between the superclusters were affected by *RUNX1::RUNX1T1* silencing. Intersection of differentially expressed genes between superclusters with those with and without *RUNX1::RUNX1T1* knockdown revealed 271 and 234 genes that are upregulated in knockdown cells and are differentially expressed between superclusters I and II, respectively (Fig. 6F). For the supercluster I/knockdown set, STRING analysis identified a cluster comprising genes related to neutrophilic granules, a ribosomal and a chaperone gene cluster (Fig. 6G, Suppl. Fig. 5C, D). In the case of the supercluster II/knockdown group, only one cluster representing genes associated with eosinophilic granules along with *CEBPE* was identified (Fig. 6H, Suppl. Fig. 5E left).

CD125, encoded by the *IL5RA* gene, is a hallmark of eosinophilic differentiation, including mature eosinophils. We recently showed that t(8;21) AML is characterized by elevated IL5RA expression and identified an IL5RA-positive leukemia stem cell (LSC) population that relies on VEGF and IL5 signaling for cell cycle entry ^23^. Consistent with this finding, high *IL5RA* transcript and CD125 surface expression were detected in both immature leukemic cell populations and more differentiated eosinophilic precursor-like ones (Suppl. Fig. 5E middle, F, G). Furthermore, we previously observed that the *IL5RA* locus contains two open chromatin regions with GATA motifs, suggesting transcriptional regulation by GATA family members^23^. Gene expression analysis comparing the two superclusters revealed significantly higher GATA2 expression in eosinophil-like cells (Suppl. Table 8, Suppl. Fig. 5E right). Cells undergoing granulocytic differentiation had lower *GATA2* expression and lacked expression of *IL5RA* (Suppl. Fig. 5E middle, right). These data show that *IL5RA* expressing t(8;21) LSCs have eosinophilic differentiation potential. Moreover, the *IL5RA* expression suggests an eosinophilic imprint of this immature cell population.

To investigate a possible imprint, we assessed the chromatin accessibility at sites associated with the differentially expressed genes from each supercluster (Fig. 6I). This result revealed that chromatin was already accessible in the PDX at sites associated with both trajectories without RUNX1::RUNX1T1 knockdown but was not accessible in CD34+ healthy peripheral blood stem cells (PBSCs) or non-t(8;21) AML cells. In aggregate, these data suggest an epigenetic imprint from RUNX1::RUNX1T1 results in a limited differentiation towards two cell fates, which progresses further following *RUXN1::RUNX1T1* silencing.

### RUNX1::RUNX1T1 knockdown depletes immature leukemic cell populations and impairs self-renewal

AML is sustained by leukemic stem cells (LSCs) that poorly respond to current treatment regimens and are a source of relapse ^73,74^. The reduction in the stem cell-specific chromatin patterns in combination with the enhanced granulocytic and eosinophilic differentiation potential prompted us to examine whether the depletion of RUNX1::RUNX1T1 eliminate LSC-enriched immature cell populations and diminished leukemic self-renewal capacity. Supporting this hypothesis, CD34-expressing leukemic cell populations exhibit a high HSC score and pseudotime analysis identified these cell populations as origin of the bidirectional differentiation (Figs. 4K, 7A, B). These populations were also lost following RUNX1::RUNX1T1 silencing and account for the overall reduction in hematopoietic stemness signatures, as indicated by GSEA (Fig. 7A–C, Suppl. Fig. 6A).

**Figure 7.**
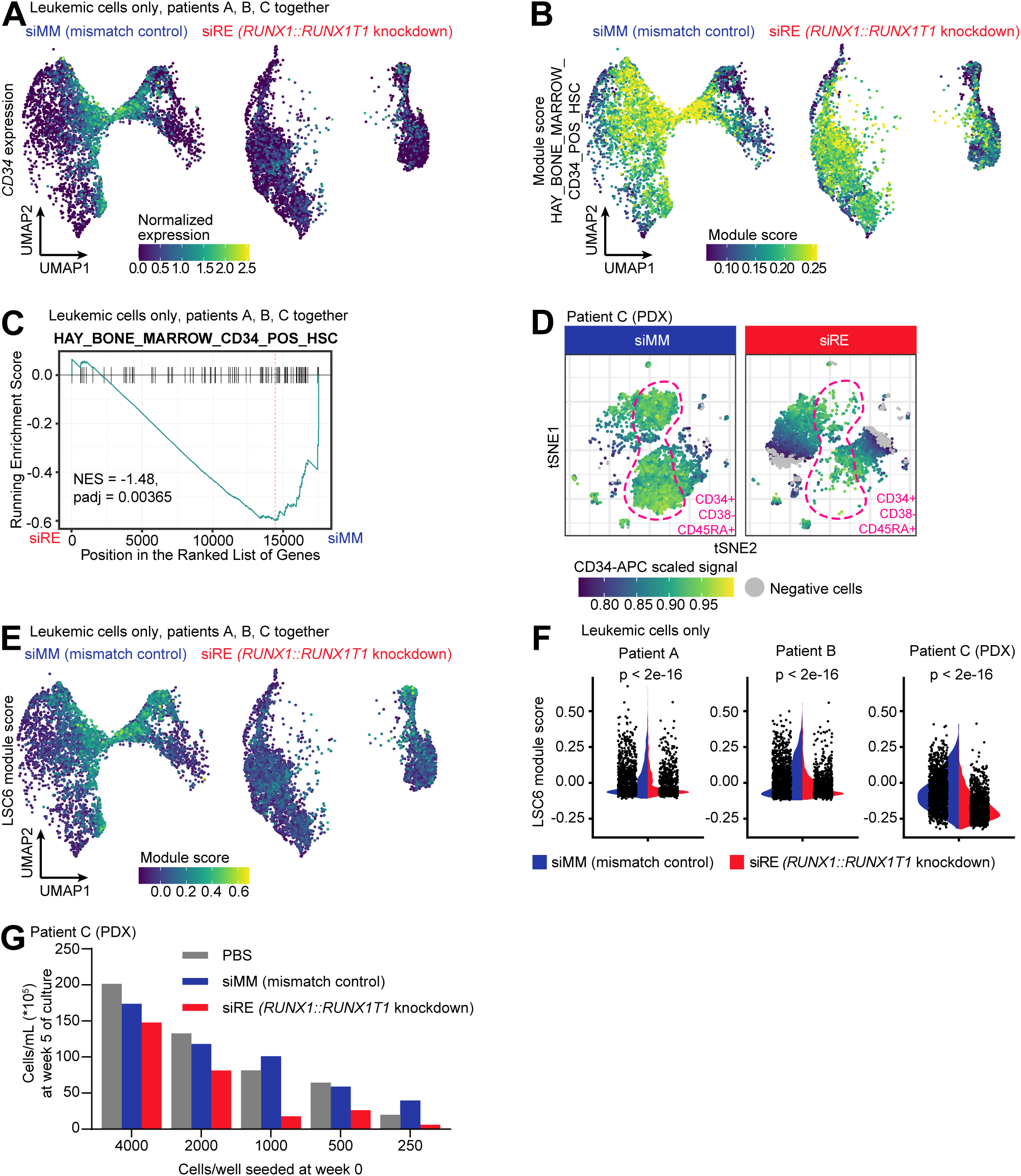
RUNX1::RUNX1T1 depletion compromises leukemic self-renewal. (A, B) UMAP plots of the integrated scRNAseq dataset of the 3 patients, split by condition (siMM- or siRE-treated), colored by normalized expression of a stemness marker *CD34* (A) and module score of the CD34-positive HSC gene set from the Human Cell Atlas bone marrow gene set collection ^55^ (B). (C) GSEA plot of the integrated scRNAseq dataset of the 3 patients, using the CD34-positive bone marrow HSC gene set ^55^. (D) tSNE plot of multi-color flow cytometry results of siRNA-treated t(8;21) AML PDX cells colored by scaled signal of CD34. CD34+CD38-CD45RA+ immature cells are highlighted with pink line. (E) UMAP plot, split by condition (siMM- or siRE-treated), colored for LSC6 gene set module score ^53^. (F) Violin plots of scRNAseq data, LSC6 ^53^ module score per siRNA-treated patient. Wilcoxon rank sum test was used to determine the p-value. (G) Bar plot depicting limiting dilution assay to evaluate reinitiation of propagation capacity of siRNA-treated PDX cells. siRE, *RUNX1::RUNX1T1* siRNA; siMM, mismatch control.

Flow cytometric analysis of eight CD markers confirmed a consistent loss of the most immature subpopulations across the PDX and all three patient samples examined (Fig. 7D, Suppl. Fig. 6B–D). Notably, patient samples exhibited heterogeneity in immature phenotypes with CD34+CD38− cells in patients B, C, and D and CD34+CD38+ cells in patient A (Suppl. Fig. 6C, D).

To specifically examine the loss of stemness in LSCs, we analyzed stemness-related transcriptional programs by assessing the LSC6 module score in relation to RUNX1::RUNX1T1 status (Fig. 7E) ^53^. *RUNX1::RUNX1T1* silencing significantly decreased the LSC6 module score across all three patient samples, with the central clusters exhibiting the highest scores (Fig. 7E, F). Furthermore, analysis of cell cycle-associated gene expression revealed that these cells preferentially reside in the G0/G1 phase, indicative of the quiescent state characteristic of LSCs ^23^(Suppl. Fig. 6E).

To functionally assess the impact of *RUNX1::RUNX1T1* depletion on leukemic self-renewal, limiting dilution assays were performed using LNP-pretreated patient-derived AML cells (Fig. 7G). *RUNX1::RUNX1T1* knockdown substantially reduced leukemic expansion capacity, confirming that silencing the fusion gene effectively diminishes the leukemia-initiating cell population.

Collectively, these findings suggest that RUNX1::RUNX1T1 maintains leukemic self-renewal potential by preventing differentiation along two distinct trajectories: granulocyte-like and aberrant eosinophil-like lineages. These findings show that therapeutic targeting of RUNX1::RUNX1T1 leads to the loss of cellular immaturity and self-renewal capacity in primary t(8;21) AML along a continuum of cell types.

## Discussion

In this study, we used an innovative siRNA delivery approach in combination with an experimentally amenable *ex vivo* culture to investigate how direct manipulation of a leukemic fusion gene affects the balance of leukemic self-renewal and differentiation in patient-derived AML cells. Here, we show that leukemic self-renewal of primary AML is strictly dependent on continuous *RUNX1::RUNX1T1* expression in agreement with previous work in cell lines ^13,15,21,75^. In addition, we demonstrate that the cellular composition of primary AML drastically changes upon *RUNX1::RUNX1T1* silencing. Knockdown of *RUNX1::RUNX1T1* by a fusion site-specific siRNA drives bidirectional granulocytic and eosinophilic differentiation at the expense of LSC-enriched immature cell populations.

The generally accepted hierarchical model of AML places LSCs as the sole cell type with self-renewal capacity and sole source of relapse on top of the hierarchy ^73,76^. This very strict organisation has been recently challenged by the observation that more mature AML cell populations can be a source of relapse by regaining an LSC-like phenotype ^77–80^. Eosinophilic cell states have also been described to regain self-renewal capability and induce relapse in a murine AML model with inducible PU.1 (also known as *SPI1*) knockdown ^81^. These findings raise the question whether *RUNX1::RUNX1T1* silencing alone is sufficient to eliminate leukemia restoring capacity or whether these fusion gene-positive descendants can lead to disease recurrence.

Our combined results reveal the existence of immature populations with granulocytic and eosinophilic potential capable of a limited degree of differentiation along these two lineages. Loss of *RUNX1::RUNX1T1* silencing leads to the almost complete elimination of the immature populations and the emergence of cells of more advanced eosinophilic and granulocytic phenotype. Notably, differentiation progressed despite the presence of UM171/UM729 and StemRegenin 1, two strong inhibitors of hematopoietic differentiation ^82,83^. Nevertheless, these inhibitors are likely to have limited the terminal differentiation of knockdown cells towards mature cell differentiation states. However, even this limited differentiation is paralleled by a substantially reduced ability of *RUNX1::RUNX1T1*-depleted cells to reinitiate leukemic proliferation under limiting dilution over a period of 5 weeks. Although we cannot formally exclude a leukemia-regenerating cell (LRC) potential for these differentiated populations, these findings strongly suggest that transient depletion of *RUNX1::RUNX1T1* impairs leukemic self-renewal in a long-lasting manner, depleting self-renewal ability along the entire continuum of leukemic cell types.

Based on recent transcriptome analyses, eosinophils, basophils and mast cells form a distinct differentiation clade separate from erythroid/megakaryocytic, granulo/monocytic and lymphoid differentiation ^54^. In contrast to the differentiation into eosinophils, the potential of t(8;21) AML to differentiate to mast cells and a link to mastocytosis is well described ^84,85^. Moreover, *RUNX1::RUNX1T1*-positive mast cells been recently identified as a source for persisting molecular measurable residual disease in disease-free patients, sometimes together with expansion of fusion gene-positive basophils ^86,87^. We have previously shown that t(8;21)-positive LSC-enriched cell populations express eosinophilic markers and employ IL5 signaling for cell cycle re-entry while preserving self-renewal capacity ^23^. Our chromatin accessibility pattern analyses indicate that these LSCs are already poised for differentiation towards the eosinophilic and granulocytic lineages. Notably, eosinophilic differentiation occurs without external addition of IL5. Since our RNA-seq data did not detect significant *IL5* transcript levels, this implies an IL5-independent induction of eosinophilic differentiation. Interestingly, C/EBPα and GATA2 cooperate in inducing eosinophilic differentiation ^88^. Furthermore, GATA2 binds to an enhancer of *CEBPE* activating its expression and may cooperate with this C/EBP member to drive expression of eosinophilic marker genes ^89^. Comparison of the expression of transcription factor genes between the granulocytic and eosinophilic superclusters revealed a similar induction of *CEBPA* expression in both superclusters upon *RUNX1::RUNX1T1* silencing, while *GATA2* and *CEBPE* expression were substantially higher in the eosinophilic supercluster. These data suggest that increasing levels of C/EBPα in conjunction with GATA2 and C/EBPε promote eosinophilic differentiation. Therefore, *RUNX1::RUNX1T1* silencing does not open new avenues of differentiation but facilitates differentiation by opening transcriptional programs along epigenetically predetermined paths.

In aggregate, we show that *RUNX1::RUNX1T1-*expressing AML cells have the intrinsic potential to differentiate along granulocytic and eosinophilic lineages. The transient silencing of the fusion gene by siRNA-loaded LNPs leads to progressed bidirectional differentiation, the elimination of LSC populations and the loss of leukemic self-renewal. These findings also warrant further research in the application of fusion gene knockdown strategies as a leukemia-specific therapeutic approach.

## Methods

### Patient characteristics

Bone marrow aspirates from AML patients from the Princess Máxima Center Biobank Facility (Project approval numbers PMC2019.059, PMC2020.184, PMCLAB2021.207) were used in this study upon signing an informed consent form by the patients or their legal representatives for use of their material for research purposes. Relevant characteristics of the patients can be found in the Suppl. Table 1.

### Patient-derived xenograft characteristics

Bone marrow aspirate from patient C was used to generate a patient-derived xenograft (PDX) as described previously ^23^. In brief, NOD.Cg-*Prkdc*^scid^ Il2rg^tm1Wjl^/SzJ (NSG) mice were injected intrafemorally with 10^6^ PDX cells. Amplified cells were harvested from the ascites and bone marrow of engrafted mice.

### LNP preparation, functionalization and characterization

LNPs were prepared as previously described ^21,24,25^. In brief, 3 volumes siRNAs dissolved in 25 mM sodium acetate (pH 4) were pumped through a microfluidic mixer (NanoAssemblr, Precision Nanosystems, Vancouver, Canada) together with 1 volume lipids dissolved in absolute ethanol at a combined flow rate of 4 ml/min. The lipid mixture contained the ionizable lipid Dlin-MC3-DMA, helper lipids cholesterol and DSPC together with the shielding lipid conjugate DMG-PEG_2000_ at a molar ratio of 50:38.5:10:1.5. Final LNP solutions were dialyzed against phosphate buffered saline (PBS). Functionalization with the Very Late Antigen-4 (VLA-4) ligand LDV conjugated to DBCO-DSPE-PEG_2000_ and characterisation were performed as prescribed previously ^24,25^.

### Cell culture

Mesenchymal stromal cells (MSCs) were obtained from healthy human bone marrow and cultivated with AML cells as previously described using either 135 nM UM171 or 1.35 μM UM729 ^24,26^. In brief, cells were seeded at a density of 7500 cells/cm^2^ one day prior to AML cells seeding to allow for expansion, with exception for the Assay for Transposase-Accessible Chromatin sequencing (ATAC-seq) and bulk RNAseq experiments. Primary patient t(8;21) leukemic bone marrow aspirate cells were co-cultured on MSCs feeders and cultivated with AML expansion medium containing peptides and cytokines ^24^. Cells were cultured at 37°C in a humidified atmosphere containing 5% CO_2_. PDX cells for single cell (sc) RNAseq and long-term culture were cultured similarly to the primary bone marrow aspirate cells, as described above, but without GM-CSF in the culture medium. PDX cells for all other experiments were cultured similarly to the primary bone marrow aspirate cells, as described above.

### LNP treatment

If not otherwise indicated, PDX and primary AML cells were seeded at a density of 10^6^ cells/ml in AML expansion medium on MSCs as described above and treated with 4 μg/ml siRNA LDV-LNPs. The next day, the cells were diluted to a density of 5 x 10^5^ cell/ml to allow expansion and collected after indicated time-points for analysis. This procedure was repeated every 72h for a total of three siRNA/vehicle doses and three days after the last dose all the groups were harvested.

### Long-term culture and limiting dilution assay

PDX cells were cultured as described above. After 72h of recovery, cells were replated on a fresh layer of MSCs and treated with 2 µg/mL of siRNA LDV-LNPs or equal vehicle volume (PBS). This procedure was repeated every 72h for a total of three siRNA/vehicle doses and three days after the last dose all the groups were harvested. Live cells were counted and five different cell densities (4000, 2000, 1000, 500 and 250 cells/well) were seeded on fresh MSCs using 96-well plates with 200 µL of AML expansion media. The cells were cultivated for 5 weeks at 37°C, 5% CO_2_ and 95% humidity with the media being replaced twice a week. Cell numbers were determined by counting with 0.04% trypan blue in a Neubauer chamber.

### LNP uptake and internal RUNX1T1 visualization

LNP uptake measurement and internal RUNX1T1 stain were done as previously described ^24^. In brief, cells were incubated for up to 24 hours with Cy3-labelled LNPs with and without LDV decoration at a siRNA concentration of 4 µg/ml. After 6 and 24 hours, cells were collected and sequentially washed in FACS buffer (0.025% m/v BSA, 0.02% m/v NaN_3_, 1% v/v FBS in PBS) and acetic acid buffer (0.5 M NaCl, 0.2 M acetic acid, pH 4), stained with antibodies and measured on the Flow cytometer. For internal RUNX1T1 staining, the cells were spun on coverslips using a Thermo Fisher Scientific Cytospin 4 Centrifuge and stained with 4’,6-diamidino-2-phenylindole (DAPI), anti-RUNX1T1 (cat #PA5-79943, Thermo Fisher Scientific, Bleiswijk, the Netherlands) and goat-anti-rabbit-IgG-AF555 (cat #405324, BioLegend, Amsterdam, the Netherlands) after permeabilization using 0.1% Triton X-100 ^24^.

### RNA extraction, cDNA synthesis and qPCR

Total RNA was extracted using the RNeasy Mini Kit (Qiagen, Hombrechtikon, Switzerland) or the NucleoSpin RNA Mini kit (Macherey-Nagel, Düren, Germany) following the corresponding supplier’s protocol. 500 ng cDNA were prepared using the RevertAid H Minus First Strand cDNA Synthesis Kit (Thermo Scientific, Bleiswijk, the Netherlands) according to manufacturer’s protocol, using the random octamer primer and the pre-incubation step of the reaction mix at 25°C for 5 min. qPCR primer sequences are provided in Suppl. Table 2. Primers for *RUNX1::RUNX1T1* were used at a concentration of 300 nM. Other primers were used at a concentration of 100 nM. cDNA samples were diluted to 15 ng input/well prior to qPCR analysis with RNAse-free H_2_O. The SsoAdvanced^TM^ Universal SYBR® Green Supermix (Bio-Rad, Veenendaal, the Netherlands) was used according to supplier’s protocol.

### Statistical analysis of qPCR results

Data were analyzed with Graphpad Prism 8 (Graphpad Software, InC., La Jolla, CA, USA) using one-tailed one-sample t-test. Differences with p-values < 0.05 were considered statistically significant.

### Protein extraction and Western blot for RUNX1::RUNX1T1

Proteins were isolated simultaneously with the RNA using the RNeasy (Qiagen, Hombrechtikon, Switzerland) or NucleoSpin RNA Mini kit (Macherey-Nagel, Düren, Germany). The column flowthrough was mixed with 2x cold acetone and incubated for 2 hours at −80°C, and dissolved in 9 M Urea buffer (4% CHAPS and 1% dithiothreitol) at a concentration of 5 x 10^4^ cell equivalents per µl. Western blotting was carried out according as previously described ^21^. Rabbit monoclonal anti-RUNX1 (1:250, #4334S, Cell Signaling Technology), mouse monoclonal anti-GAPDH (1:10000, #AM4300, Invitrogen, Thermo Fisher Scientific, Bleiswijk, the Netherlands) were used as primary antibodies. Goat anti-mouse (1:10000, #P0447, Agilent, Abcoude, the Netherlands) or anti-rabbit (1:10000, #sc-2004, Santa Cruz Biotechnology, Heidelberg, Germany) polyclonal IgG HRP-conjugates were used as secondary antibodies.

### Bulk RNA-seq

Cells for bulk RNAseq were harvested 3 days after a single LDV-LNP treatment. Total RNA was extracted using the RNeasy Mini Kit (Qiagen) or the NucleoSpin RNA Mini kit (Macherey-Nagel,), following the corresponding supplier’s protocol. Quality control of the extracted total RNA was done on the Bioanalyzer instrument using the RNA 6000 Nano Kit (Agilent). 500 ng of RNA, with RNA integrity number larger than 9, was provided to the Diagnostic department of the Princess Máxima Center for Pediatric Oncology, who prepared the libraries using the KAPA RNA HyperPrep Kit with RiboErase (Roche, Woerden, the Netherlands) and sequenced on a NovaSeq 6000 using the S4 Reagent Kit v1.5 300 cycles (Illumina, Eindhoven, the Netherlands) with the 2×150bp setup (paired-end sequencing).

### Bulk ATAC-seq

Cells for the bulk ATACseq were harvested 3 days after a single siRNA LDV-LNP treatment. ATAC-seq was performed according to the Omni-ATAC protocol as previously described ^27,28^. Briefly, 5 x 10^4^ cells were treated with Miltenyi Dead Cell Removal Kit following supplier’s protocol (Miltenyi Biotec, Leiden, the Netherlands) and consequently treated with Tagment DNA TDE1 Enzyme (Illumina) and subjected to library preparation using the NEBNext High-Fidelity 1x PCR Master Mix (New England Biolabs, Ipswich, MA, USA) with barcoded PCR primers (custom-synthesized by Integrated DNA Technologies; sequences as described ^29^) followed by paired-end 150 bp sequencing on a NovaSeq 6000 (Illumina). Samples were sequenced by Novogene (Cambridge, United Kingdom).

### scRNA-seq

Cells for scRNA-seq were harvested 3 days after 2 consecutive siRNA LDV-LNP treatments with 3 days in between treatments. 3’ scRNA library preparation was done using the 10X Genomics v3.1 chemistry (10X Genomics protocol CG000206 Rev D, 10X Genomics, Leiden, the Netherlands), preceded by Cell Hashing ^30^. Briefly, after harvesting, cells were washed with a sterile solution of 1% m/v BSA in DPBS (#14190144, Gibco – Thermo Fisher Scientific), blocked with 20 µg/ml human IgG for 10 min on ice and stained with 2 µg of a TotalSeq™-B Hashtag Antibody pool as described by the provider (Biolegend, Amsterdam, the Netherlands) in 50 ul of 1% m/v BSA in DPBS (#14190144, Gibco – Thermo Fisher Scientific) for 20 min on ice. The list of the Hashtag antibodies used can be found in Supplementary Table 3.

Hashed cells were washed 3 times with 1% m/v BSA in DPBS (#14190144, Gibco – Thermo Fisher Scientific) and resuspended in 1% m/v BSA in DPBS. Quantity, viability and single cellularity of the hashed cells were assessed visually under a microscope using 1:1 staining of the cell suspension with 0.04% Trypan Blue and a Neubauer chamber.

Hashed cells were pooled pairwise: from each patient, cells treated with either *RUNX1::RUNX1T1* siRNA LDV-LNP or control mismatch siRNA LDV-LNP were pooled. 12,000 cells per sample (24000 cells per sample pool) for the primary patient samples and 10,000 cells per sample (20000 cells per sample pool) for the PDX were loaded onto the Chromium Next GEM Chip G (10X Genomics, Leiden, the Netherlands). Subsequently, 3’ scRNA library and Hashtag Oligo (HTO) libraries were prepared following the 10X Genomics protocol CG000206 Rev D using 11 PCR cycles of cDNA amplification for the primary patient samples and 12 cycles for the PDX. cDNA quality control and yield assessment were done on the Bioanalyzer instrument using the High Sensitivity DNA Kit (Agilent). Based on the cDNA yield, we performed 12 cycles of sample index (SI) PCR for primary patient samples and 11 cycles of SI-PCR for the PDX. Concentration of the libraries for loading in the sequencer was measured on the Qubit instrument with the dsDNA Quantitation High Sensitivity reagent kit (#Q32854, Thermo Fisher Scientific).

3’scRNA and HTO libraries were sequenced on a NovaSeq 6000 (Illumina) sequencer with the read setup indicated by the manufacturer (10X Genomics protocol CG000206 Rev D) at a targeted depth of ≥25000 read pairs per cell for the 3’ scRNA libraries and ≥2400 read pairs per cell for the HTO libraries.

### Flow cytometry

1–2 x 10^5^ cells per sample were stained in V-bottom 96-well plates. Cells were washed twice with FACS buffer followed by staining for 45 min on ice with the antibody cocktail for immunophenotyping (Suppl. Tables 4, 5), 20 µg/ml human IgG and the viability dye solution: for primary AML patient samples (patients A, B, D) this was 1 ul of the working solution of ViaKrome 808 Fixable Viability Dye (#C36628, Beckman Coulter, Woerden, the Netherlands) in 50 µl BD Horizon™ Brilliant Stain Buffer (#563794, BD Biosciences, Drachten, the Netherlands); for the PDX (patient C) it was 0.5 ul of the working solution of the Zombie UV (#423108, Biolegend, Amsterdam, the Netherlands) in 50 ul FACS buffer. Cells were then washed twice with FACS buffer and fixed in 2% paraformaldehyde (PFA) in PBS overnight at 4°C. Samples were stored in the dark at 4 °C for up to 7 days and their fluorescence was recorded on a Cytoflex LX cytometer (Beckman Coulter). Compensation was recorded from single stains of UltraComp eBeads™ Compensation Beads (#01-2222-42, Invitrogen, Thermo Fisher Scientific) with the fluorescently labelled antibodies or from a 3:3:3:1 mix of ME-1, Kasumi-1, SEM cell lines and MSC for the unstained sample and the live/dead single stain.

### Bulk RNAseq data analysis

The gene counts were produced by the Big Data Core of the Princess Máxima Center for Pediatric Oncology by aligning the sequencing results to the GRCh38 reference human genome with STAR v2.7.2d ^31^. Differential expression (DE) analysis was done in R with DESeq2 package ^32^ using standard parameters. GO enrichment analysis was performed with the enrichGO function with a p-value cutoff of 0.05 and GSEA was performed with the GSEA function of the clusterProfiler library ^33^. Results were visualised using the BiocGenerics ^34^, circlize ^35^, ComplexHeatmap ^36^, EnhancedVolcano ^37^ and enrichplot ^38^ packages. Bigwig files were made using the bamCoverage function in deepTools 3.5.0 ^39^. The workflow can be followed in detail on Github (see “Code availability” section).

### ATAC-seq data analysis

ATAC fastq files were processed with Trimmomatic v0.39 to remove sequencing adaptors and low-quality reads. Trimmed reads were aligned to the human genome (version hg38) using Bowtie2 v2.4.4 ^40^ using the setting --very-sensitive-local. PCR duplicates were removed using the MarkDuplicates function in Picard 2.21.1. Bigwig files were made using the bamCoverage function in deepTools 3.5.0 ^39^ and normalized as counts per million (cpm). Peaks were called using MACS2 v2.2.7.1^41^. using the settings -q 0.0005 -B --trackline --nomodel --shift −100 --extsize 200 and peaks were filtered against the ENCODE blacklist.

To assess replicates, a peak union was generated from all peaks using bedtools v2.29.2 ^42^ merge. The average tag-density in a 400-bp window centered on the peak union summits was calculated for each sample using the annotatePeaks.pl function in Homer v4.11 ^43^ using the bedGraph files generated by MACS2. Tag counts were CPM normalised and distal sites only selected (>1.5kb from annotated TSS). Tag counts had a pseudocount of 0.1 added and were log2 transformed, and Pearson correlation calculated.

To carry out differential chromatin accessibility analysis a peak union was generated from only peaks shared in all 3 replicates using bedtools intersect and merge. Tag counts in these peaks were obtained using featureCounts ^44^ from the Subread package v2.0.1 using the option -p. Differentially accessible peaks were calculated with DESeq2^32^, peaks were considered differential if p<0.05, fold change >2. De novo motif discovery was carried out in the differential peaks using HOMER findMotifsGenome -noknown. Density plots were generated using Homer annotatePeaks.pl function using the bedGraph files generated by MACS2 or motif position weight matrix files built into HOMER, with the options -size 2000 -hist 10 -ghist and plotted order by fold change using JavaTreeView 1.1.6.

Comparison with healthy cells used data from GSE74912 ^45^ and GSE150023^46^, a peak set specific to each cell type was calculated and this was compared to the fold change-ordered chromatin accessibility to determine if the cell type specific peaks were present or absent, this binary was plotted with JavaTreeView. Comparison with non-t(8;21) and PBSCs used data from GSE108316 ^47^, samples ITD-1, ITD-2 and PBSC-2. Average profiles were created using CPM normalized bigwig files using the computeMatrix, plotProfile functions in deepTools, plotting the output in R or deepTools plotHeatmap function. For CEBPE distal site the bed file annotated contained the genomic coordinates derived from the peak union; sites associated with Supercluster markers were obtained from the peak union annotated to the nearest gene with HOMER.

Bigwig files were visualised using the UCSC genome browser, alongside DNaseI-seq d2 knockdown and RUNX1::RUNX1T1 ChIP-seq from Kasumi-1 obtained from GSE29225 ^48^). UCSC screenshots were made as described previously from the UCSC Genome Browser ^14,49^.

The workflow can be followed in detail on Github (see “Code availability” section).

### Flow cytometry data analysis

Compensation was applied automatically during the acquisition by the CytExpert software (Beckman Coulter, Brea, CA, USA). The resulting datasets were gated manually in FlowJo v10.9.0 software (BD Biosciences, Ashland, OR, USA): first, for live singlet cells; the results for primary patient samples (patients A, B, D) were additionally gated for CD73-negative cells to exclude MSCs; on these populations, positivity gates for all the remaining fluorescent markers were set manually based on the Fluorescence Minus One (FMO) or unstained controls. The data for live singlet cells with the positivity thresholds for all other fluorescent markers were taken for further analysis in R. Dimensionality reduction was done in R v4.0.5 using a compute node of the Utrecht HPC running on Rocky Linux 8.9. The entire workflow can be followed on Github (see “Code availability” section). Briefly, the FCS files were read with the flowCore package v2.2.0 (Ellis et al. 2023) and their metadata such as patient IDs and conditions pre-entered manually to an Excel sheet were read with the readxl package (part of the tidyverse super-package v1.3.2). Inverse hyperbolic sine transformation was applied to the fluorescent marker area intensities, and the transformed values were scaled to a scale from 0 to 1. Transformed and scaled area intensity values of the fluorescent markers of 5000 random cells from each condition were taken for a principal component analysis (PCA) which was performed using the FactoMineR package v2.6, and for t-distributed stochastic neighbor embedding (tSNE) which was performed using the Rtsne package v0.16. The cells were then visualized in the resulting tSNE coordinates using the ggplot2 v3.3.6 package.

### scRNA-seq data analysis

Base calls were automatically converted to the FASTQ format with the software on the sequencer. All downstream analysis was done on compute nodes of the Utrecht High Performance Compute facility (Utrecht HPC) running under Rocky Linux 8.8 or 8.9. The FASTQ reads were aligned onto the reference transcriptome (GRCh38–2020–A, https://cf.10xgenomics.com/supp/cell-exp/refdata-gex-GRCh38-2020-A.tar.gz, accession date 2025-02-10) and the reference HTO sequences using the Cell Ranger v7.1.0 software (10X Genomics). The resulting Binary Alignment Map (BAM) files were used as input for the Souporcell software ^50^ to demultiplex the datasets based on the donor-derived Single Nucleotide Polymorphisms (SNPs). The gene and HTO read counts from Cell Ranger, and donor information from Souporcell were further analysed in R v4.3.2. The workflow can be followed in detail on Github (see “Code availability” section).

Briefly, the counts were loaded into the Seurat v4.4.0 package ^51^. The experimental conditions were demultiplexed with the HTODemux function with the positive quantile set at 0.95. The donors were demultiplexed based on the Souporcell assignment. HTO-based and donor-based multiplets were excluded from further analysis, no further doublet filtering was done. Cells with ≥7% of mitochondrion-encoded transcripts and transcripts of ≤750 genes were excluded from further analysis. Data transformation was performed with the SCTransform function, with regression for the differences between G2M and S-phase-associated transcripts. Principal component analysis (PCA) was done with the RunPCA function, and the suitable number of principal components (PCs) for further analysis was determined using elbow plots of variance per each PC. Neighbors and Uniform Manifold Approximation and Projection (UMAP) coordinates were found with the FindNeighbors and RunUMAP functions, respectively. Mesenchymal stromal cells (MSCs) that had served as feeder cells in the cocultures were identified with the help of Souporcell output and MSC marker gene (*NT5E, THY1*) expression, and excluded from further analysis. In the remaining cells, clusters were found with the Louvain algorithm embedded into the FindClusters function, with the resolution parameter set at 0.25. Distinction of clusters with malignant and non-malignant cells was achieved by plotting the *RUNX1T1* expression density using the Nebulosa package v1.10.0 ^52^. Clusters with low expression density of *RUNX1T1* were considered non-malignant and were further analysed separately. Differentially expressed genes (DEGs) were determined using the FindMarkers function of the Seurat package. We visualized the DEGs of the non-malignant AML patient cells with the EnhancedVolcano package v1.20.0 ^37^. GSEA was done with the GSEA function of the clusterProfiler library ^37^, using log2 fold changes of genes that were expressed in >1% of the cells in each dataset. Pediatric leukemic stem cell (LSC6) score was calculated for leukemic cells using the ModuleScore function of the Seurat package to account for the sparcity of the scRNAseq data ^53^.

Cell type predictions for the leukemic cells from each patient separately, using the Zeng reference dataset were made through the projection algorithm within the Seurat package (FindTransferAnchors and TransferData functions) ^54^. We integrated the leukemic cells from different patients using Seurat’s SCTIntegration approach. Pseudotime analysis was done with the monocle3 package v1.0.0 (Qiu et al. 2017; Cao et al. 2019). Cells with the earliest pseudotime were assigned based on the highest module score of the HAY_BONE_MARROW_CD34_POS_HSC gene set from MSigDB (www.gsea-msigdb.org/, accession date 2025-02-10, ^55^). Clustering of the integrated object was done with Seurat’s FindClusters function implementing the Leiden algorithm. Marker genes of the clusters were found using the FindAllMarkers function. Differentially expressed genes in different experimental conditions and clusters were visualized with the nVennR package^56^. Protein network analyses were performed using STRING version 12 (version11.string-db.org<u>, accession date 2025-02-10</u>).

## Data & code availability

The raw sequencing data generated in this work have been uploaded to the European genome-phenome archive under the accession number (submission in progress). For patient privacy protection, the access is controlled by the Data Access Committee (DAC) of the Princess Máxima Center for Pediatric Oncology. Researchers can obtain access by submitting a project proposal to the DAC. The reference healthy bone marrow dataset was kindly provided by Andy Zeng ^54^. Bulk RNAseq counts of Kasumi-1 and SKNO-1 cell lines upon RUNX1::RUNX1T1 depletion ^21^ are available at Gene Expression Omnibus (www.ncbi.nlm.nih.gov/geo) under entry GSE217113. The processed data generated in this work, namely the compensated FCS files from flow cytometry, gene counts for bulk RNAseq, bedGraph files for bulk ATACseq and Seurat objects for scRNAseq, have been made publicly available at Zenodo (doi.org/10.5281/zenodo.14578307). Code is available at https://github.com/polinaderev/runx1-runx1t1-kd-aml.

## Acknowledgements

This work was supported by KiKa program grant 329 and KiKa project grant 460 to O.H. Work in C.Bonifer’s lab was funded by grants from UKRI ((MR/S021469/1) and Blood Cancer UK (15001). C.B. is currently funded by the Novo Nordisk Foundation Center for Stem Cell medicine, reNEW, (Novo Nordisk Foundation grant number NNF21CC0073729). SK was supported by a Leukaemia UK John Goldman Fellowship (2023/JGF/004) and Blood Cancer UK (20006). The sponsors of this study are public or nonprofit organizations that support science in general. They had no role in gathering, analyzing, or interpreting the data. We thank the Diagnostics facility and Big Data Core of the Princess Máxima Center for performing RNA-seq and supporting data processing and analysis. We also thank Zhijun Yu, Peter Brazda, Elizabeth Schweighart (all employees of the Princess Máxima Center at the time of the experiments) and the Single Cell Genomics facility of the Princess Máxima Center for help with scRNA-seq experiments. We thank Sebastian Ramisch (intern at the Princess Máxima Center at the time of the experiments) for the help with the preparation of primary AML cells for flow cytometry. We thank the Big Data Core of the Princess Máxima Center for preparing bulk RNAseq libraries, sequencing them, performing their alignment and guiding us through the EGA submission. We are grateful to all members of the Heidenreich–Vormoor group took part in the discussions about the research. We thank the patients and their families for allowing to use the patients’ samples for the research.

## Author contribution and conflicts of interest

The study was conceived by P.K.D., L.E.S. and O.H. Experimental design was performed by L.E.S., P.K.D., L.D.M.C. Experimental work was performed by L.E.S. (everything when not otherwise stated), L.D.M.C. (LNP-siRNA preparation for some experiments, long-term culture and limiting dilution assay), P.K.D. (flow cytometry of primary AML samples, scRNAseq library preparation), L.vdB. (flow cytometry of primary AML samples), A.T.vO. (Western blotting), A.K.-H. (optimization of AML-MSC co-culture conditions). Data was analyzed by P.K.D. (bulk RNAseq, scRNAseq, flow cytometry), S.G.K. (ATACseq), M.A. (bulk RNAseq) and L.E.S. (qPCR, flow cytometry). L.E.S., P.K.D., O.H., S.G.K. and L.D.M.C. wrote the original draft. P.K.D., L.E.S., S.G.K. and O.H. prepared the figures. O.H. acquired the funding. O.H., H.J.V., R.S., C.B. and C.M.Z. supervised the work. All authors contributed to the review and editing of the manuscript.

All authors declared that there is no relevant conflict of interest.

## References

1 Elgarten, C. W. & Aplenc, R. Pediatric acute myeloid leukemia: updates on biology, risk stratification, and therapy. Curr Opin Pediatr 32, 57–66 (2020). 10.1097/MOP.0000000000000855

2 Dohner, H. et al. Diagnosis and management of AML in adults: 2022 recommendations from an international expert panel on behalf of the ELN. Blood 140, 1345–1377 (2022). 10.1182/blood.2022016867

3 Egan, G., Chopra, Y., Mourad, S., Chiang, K. Y. & Hitzler, J. Treatment of acute myeloid leukemia in children: A practical perspective. Pediatr Blood Cancer 68, e28979 (2021). 10.1002/pbc.28979

4 Lim, J. J. et al. Time independent factors that predict relapse in adults with acute myeloid leukemia. Blood Cancer J 14, 5 (2024). 10.1038/s41408-023-00954-z

5 Zarnegar-Lumley, S., Caldwell, K. J. & Rubnitz, J. E. Relapsed acute myeloid leukemia in children and adolescents: current treatment options and future strategies. Leukemia 36, 1951–1960 (2022). 10.1038/s41375-022-01619-9

6 Rasche, M., et al. Survival Following Relapse in Children with Acute Myeloid Leukemia: A Report from AML-BFM and COG. Cancers (Basel) 13 (2021). 10.3390/cancers13102336

7 De Kouchkovsky, I. & Abdul-Hay, M. ‘Acute myeloid leukemia: a comprehensive review and 2016 update’. Blood Cancer J 6, e441 (2016). 10.1038/bcj.2016.50

8 Martens, J. H. & Stunnenberg, H. G. The molecular signature of oncofusion proteins in acute myeloid leukemia. FEBS Lett 584, 2662–2669 (2010). 10.1016/j.febslet.2010.04.002

9 Creutzig, U. et al. Changes in cytogenetics and molecular genetics in acute myeloid leukemia from childhood to adult age groups. Cancer 122, 3821–3830 (2016). 10.1002/cncr.30220

10 Borkhardt, A. & Heidenreich, O. RNA interference as a potential tool in the treatment of leukaemia. Expert Opin Biol Ther 4, 1921–1929 (2004). 10.1517/14712598.4.12.1921

11 Schmid, H., Jaeger, B. A., Lohse, J. & Suttorp, M. Longitudinal growth retardation in a prepuberal girl with chronic myeloid leukemia on long-term treatment with imatinib. Haematologica 94, 1177–1179 (2009). 10.3324/haematol.2009.008359

12 Miyoshi, H. et al. The t(8;21) translocation in acute myeloid leukemia results in production of an AML1-MTG8 fusion transcript. EMBO J 12, 2715–2721 (1993).

13 Heidenreich, O. et al. AML1/MTG8 oncogene suppression by small interfering RNAs supports myeloid differentiation of t(8;21)-positive leukemic cells. Blood 101, 3157–3163 (2003). 10.1182/blood-2002-05-1589

14 Martinez-Soria, N. et al. The Oncogenic Transcription Factor RUNX1/ETO Corrupts Cell Cycle Regulation to Drive Leukemic Transformation. Cancer Cell 34, 626–642 e628 (2018). 10.1016/j.ccell.2018.08.015

15 Ptasinska, A. et al. Identification of a dynamic core transcriptional network in t(8;21) AML that regulates differentiation block and self-renewal. Cell Rep 8, 1974–1988 (2014). 10.1016/j.celrep.2014.08.024

16 Gardini, A. et al. AML1/ETO oncoprotein is directed to AML1 binding regions and co-localizes with AML1 and HEB on its targets. PLoS Genet 4, e1000275 (2008). 10.1371/journal.pgen.1000275

17 Thomas, M. et al. Targeting MLL-AF4 with short interfering RNAs inhibits clonogenicity and engraftment of t(4;11)-positive human leukemic cells. Blood 106, 3559–3566 (2005). 10.1182/blood-2005-03-1283

18 Scherr, M. et al. Specific inhibition of bcr-abl gene expression by small interfering RNA. Blood 101, 1566–1569 (2003). 10.1182/blood-2002-06-1685

19 Manara, E. et al. MLL-AF6 fusion oncogene sequesters AF6 into the nucleus to trigger RAS activation in myeloid leukemia. Blood 124, 263–272 (2014). 10.1182/blood-2013-09-525741

20 Jyotsana, N. et al. RNA interference efficiently targets human leukemia driven by a fusion oncogene in vivo. Leukemia 32, 224–226 (2018). 10.1038/leu.2017.269

21 Issa, H. et al. Nanoparticle-mediated targeting of the fusion gene RUNX1/ETO in t(8;21)-positive acute myeloid leukaemia. Leukemia 37, 820–834 (2023). 10.1038/s41375-023-01854-8

22 Ansari, A. S. et al. Lipopolymer mediated siRNA delivery targeting aberrant oncogenes for effective therapy of myeloid leukemia in preclinical animal models. J Control Release (2024). 10.1016/j.jconrel.2024.02.018

23 Kellaway, S. G. et al. Leukemic stem cells activate lineage inappropriate signalling pathways to promote their growth. Nat Commun 15, 1359 (2024). 10.1038/s41467-024-45691-4

24 Swart, L. E. et al. Increased Bone Marrow Uptake and Accumulation of Very-Late Antigen-4 Targeted Lipid Nanoparticles. Pharmaceutics 15 (2023). 10.3390/pharmaceutics15061603

25 Swart, L. E. et al. A robust post-insertion method for the preparation of targeted siRNA LNPs. Int J Pharm 620, 121741 (2022). 10.1016/j.ijpharm.2022.121741

26 Pal, D. et al. hiPSC-derived bone marrow milieu identifies a clinically actionable driver of niche-mediated treatment resistance in leukemia. Cell Rep Med 3, 100717 (2022). 10.1016/j.xcrm.2022.100717

27 Corces, M. R. et al. An improved ATAC-seq protocol reduces background and enables interrogation of frozen tissues. Nat Methods 14, 959–962 (2017). 10.1038/nmeth.4396

28 Buenrostro, J. D., Wu, B., Chang, H. Y. & Greenleaf, W. J. ATAC-seq: A Method for Assaying Chromatin Accessibility Genome-Wide. Curr Protoc Mol Biol 109, 21 29 21–21 29 29 (2015). 10.1002/0471142727.mb2129s109

29 Buenrostro, J. D., Giresi, P. G., Zaba, L. C., Chang, H. Y. & Greenleaf, W. J. Transposition of native chromatin for fast and sensitive epigenomic profiling of open chromatin, DNA-binding proteins and nucleosome position. Nat Methods 10, 1213–1218 (2013). 10.1038/nmeth.2688

30 Stoeckius, M. et al. Cell Hashing with barcoded antibodies enables multiplexing and doublet detection for single cell genomics. Genome Biol 19, 224 (2018). 10.1186/s13059-018-1603-1

31 Dobin, A. et al. STAR: ultrafast universal RNA-seq aligner. Bioinformatics 29, 15–21 (2013). 10.1093/bioinformatics/bts635

32 Love, M. I., Huber, W. & Anders, S. Moderated estimation of fold change and dispersion for RNA-seq data with DESeq2. Genome biology 15, 550 (2014).

33 Wu, T. et al. clusterProfiler 4.0: A universal enrichment tool for interpreting omics data. Innovation (Camb) 2, 100141 (2021). 10.1016/j.xinn.2021.100141

34 Huber, W. et al. Orchestrating high-throughput genomic analysis with Bioconductor. Nat Methods 12, 115–121 (2015). 10.1038/nmeth.3252

35 Gu, Z., Gu, L., Eils, R., Schlesner, M. & Brors, B. circlize Implements and enhances circular visualization in R. Bioinformatics 30, 2811–2812 (2014). 10.1093/bioinformatics/btu393

36 Gu, Z., Eils, R. & Schlesner, M. Complex heatmaps reveal patterns and correlations in multidimensional genomic data. Bioinformatics 32, 2847–2849 (2016). 10.1093/bioinformatics/btw313

37 Bilghe, K., Rana, S. & Lewis, M. EnhancedVolcano: Publication-ready volcano plots with enhanced colouring and labeling. . R package 1.24.0 (2024). 10.18129/B9.bioc.EnhancedVolcano

38 Yu, G. enrichplot: Visualization of Functional Enrichment Result. . R package 1.27.4 (2025). 10.18129/B9.bioc.enrichplot

39 Ramirez, F. et al. deepTools2: a next generation web server for deep-sequencing data analysis. Nucleic Acids Res 44, W160–165 (2016). 10.1093/nar/gkw257

40 Langmead, B. & Salzberg, S. L. Fast gapped-read alignment with Bowtie 2. Nat Methods 9, 357–359 (2012). 10.1038/nmeth.1923

41 Zhang, Y. et al. Model-based analysis of ChIP-Seq (MACS). Genome Biol 9, R137 (2008). 10.1186/gb-2008-9-9-r137

42 Quinlan, A. R. & Hall, I. M. BEDTools: a flexible suite of utilities for comparing genomic features. Bioinformatics 26, 841–842 (2010). 10.1093/bioinformatics/btq033

43 Heinz, S. et al. Simple combinations of lineage-determining transcription factors prime cis-regulatory elements required for macrophage and B cell identities. Mol Cell 38, 576–589 (2010). 10.1016/j.molcel.2010.05.004

44 Liao, Y., Smyth, G. K. & Shi, W. featureCounts: an efficient general purpose program for assigning sequence reads to genomic features. Bioinformatics 30, 923–930 (2014). 10.1093/bioinformatics/btt656

45 Corces, M. R. et al. Lineage-specific and single-cell chromatin accessibility charts human hematopoiesis and leukemia evolution. Nat Genet 48, 1193–1203 (2016). 10.1038/ng.3646

46 Perez, C. et al. Immunogenomic identification and characterization of granulocytic myeloid-derived suppressor cells in multiple myeloma. Blood 136, 199–209 (2020). 10.1182/blood.2019004537

47 Assi, S. A. et al. Subtype-specific regulatory network rewiring in acute myeloid leukemia. Nat Genet 51, 151–162 (2019). 10.1038/s41588-018-0270-1

48 Ptasinska, A. et al. Depletion of RUNX1/ETO in t(8;21) AML cells leads to genome-wide changes in chromatin structure and transcription factor binding. Leukemia 26, 1829–1841 (2012). 10.1038/leu.2012.49

49 Perez, G. et al. The UCSC Genome Browser database: 2025 update. Nucleic Acids Res 53, D1243–D1249 (2025). 10.1093/nar/gkae974

50 Heaton, H. et al. Souporcell: robust clustering of single-cell RNA-seq data by genotype without reference genotypes. Nat Methods 17, 615–620 (2020). 10.1038/s41592-020-0820-1

51 Hao, Y. et al. Integrated analysis of multimodal single-cell data. Cell 184, 3573–3587 e3529 (2021). 10.1016/j.cell.2021.04.048

52 Alquicira-Hernandez, J. & Powell, J. E. Nebulosa recovers single-cell gene expression signals by kernel density estimation. Bioinformatics 37, 2485–2487 (2021). 10.1093/bioinformatics/btab003

53 Elsayed, A. H. et al. A six-gene leukemic stem cell score identifies high risk pediatric acute myeloid leukemia. Leukemia 34, 735–745 (2020). 10.1038/s41375-019-0604-8

54 Zeng, A. G. X. et al. Precise single-cell transcriptomic mapping of normal and leukemic cell states reveals unconventional lineage priming in acute myeloid leukemia. bioRxiv (2023). 10.1101/2023.12.26.573390

55 Hay, S. B., Ferchen, K., Chetal, K., Grimes, H. L. & Salomonis, N. The Human Cell Atlas bone marrow single-cell interactive web portal. Exp Hematol 68, 51–61 (2018). 10.1016/j.exphem.2018.09.004

56 Perez-Silva, J. G., Araujo-Voces, M. & Quesada, V. nVenn: generalized, quasi-proportional Venn and Euler diagrams. Bioinformatics 34, 2322–2324 (2018). 10.1093/bioinformatics/bty109

57 Chigaev, A. et al. Real time analysis of the affinity regulation of alpha 4-integrin. The physiologically activated receptor is intermediate in affinity between resting and Mn(2+) or antibody activation. J Biol Chem 276, 48670–48678 (2001). 10.1074/jbc.M103194200

58 Cai, D. H. et al. C/EBP alpha:AP-1 leucine zipper heterodimers bind novel DNA elements, activate the PU.1 promoter and direct monocyte lineage commitment more potently than C/EBP alpha homodimers or AP-1. Oncogene 27, 2772–2779 (2008). 10.1038/sj.onc.1210940

59 Friedman, A. D. C/EBPalpha in normal and malignant myelopoiesis. Int J Hematol 101, 330–341 (2015). 10.1007/s12185-015-1764-6

60 Obier, N. & Bonifer, C. Chromatin programming by developmentally regulated transcription factors: lessons from the study of haematopoietic stem cell specification and differentiation. FEBS Lett 590, 4105–4115 (2016). 10.1002/1873-3468.12343

61 Avellino, R. & Delwel, R. Expression and regulation of C/EBPalpha in normal myelopoiesis and in malignant transformation. Blood 129, 2083–2091 (2017). 10.1182/blood-2016-09-687822

62 Yamanaka, R. et al. CCAAT/enhancer binding protein epsilon is preferentially up-regulated during granulocytic differentiation and its functional versatility is determined by alternative use of promoters and differential splicing. Proc Natl Acad Sci U S A 94, 6462–6467 (1997). 10.1073/pnas.94.12.6462

63 Keeshan, K., Santilli, G., Corradini, F., Perrotti, D. & Calabretta, B. Transcription activation function of C/EBPalpha is required for induction of granulocytic differentiation. Blood 102, 1267–1275 (2003). 10.1182/blood-2003-02-0477

64 Pabst, T. et al. AML1-ETO downregulates the granulocytic differentiation factor C/EBPalpha in t(8;21) myeloid leukemia. Nat Med 7, 444–451 (2001). 10.1038/86515

65 Shyamsunder, P. et al. Identification of a novel enhancer of CEBPE essential for granulocytic differentiation. Blood 133, 2507–2517 (2019). 10.1182/blood.2018886077

66 Theilgaard-Monch, K. et al. Transcription factor-driven coordination of cell cycle exit and lineage-specification in vivo during granulocytic differentiation : In memoriam Professor Niels Borregaard. Nat Commun 13, 3595 (2022). 10.1038/s41467-022-31332-1

67 Tonks, A. et al. Transcriptional dysregulation mediated by RUNX1-RUNX1T1 in normal human progenitor cells and in acute myeloid leukaemia. Leukemia 21, 2495–2505 (2007). 10.1038/sj.leu.2404961

68 Dunne, J. et al. siRNA-mediated AML1/MTG8 depletion affects differentiation and proliferation-associated gene expression in t(8;21)-positive cell lines and primary AML blasts. Oncogene 25, 6067–6078 (2006). 10.1038/sj.onc.1209638

69 Schoenherr, C. et al. Stable depletion of RUNX1-ETO in Kasumi-1 cells induces expression and enhanced proteolytic activity of Cathepsin G and Neutrophil Elastase. PLoS One 14, e0225977 (2019). 10.1371/journal.pone.0225977

70 ElTanbouly, M. A., Croteau, W., Noelle, R. J. & Lines, J. L. VISTA: a novel immunotherapy target for normalizing innate and adaptive immunity. Semin Immunol 42, 101308 (2019). 10.1016/j.smim.2019.101308

71 Elghetany, M. T. & Patel, J. Assessment of CD24 expression on bone marrow neutrophilic granulocytes: CD24 is a marker for the myelocytic stage of development. Am J Hematol 71, 348–349 (2002). 10.1002/ajh.10176

72 Barkal, A. A. et al. CD24 signalling through macrophage Siglec-10 is a target for cancer immunotherapy. Nature 572, 392–396 (2019). 10.1038/s41586-019-1456-0

73 Reya, T., Morrison, S. J., Clarke, M. F. & Weissman, I. L. Stem cells, cancer, and cancer stem cells. Nature 414, 105–111 (2001). 10.1038/35102167

74 Essers, M. A. & Trumpp, A. Targeting leukemic stem cells by breaking their dormancy. Mol Oncol 4, 443–450 (2010). 10.1016/j.molonc.2010.06.001

75 Stengel, K. R., Ellis, J. D., Spielman, C. L., Bomber, M. L. & Hiebert, S. W. Definition of a small core transcriptional circuit regulated by AML1-ETO. Molecular Cell (2020). 10.1016/j.molcel.2020.12.005

76 Bonnet, D. & Dick, J. Human acute myeloid leukemia is organized as a hierarchy that originates from a primitive hematopoietic cell. (1997).

77 Boyd, A. L. et al. Identification of Chemotherapy-Induced Leukemic-Regenerating Cells Reveals a Transient Vulnerability of Human AML Recurrence. Cancer Cell 34, 483–498 e485 (2018). 10.1016/j.ccell.2018.08.007

78 McKenzie, M. D. et al. Interconversion between Tumorigenic and Differentiated States in Acute Myeloid Leukemia. Cell Stem Cell 25, 258–272 e259 (2019). 10.1016/j.stem.2019.07.001

79 Pei, S. et al. Monocytic Subclones Confer Resistance to Venetoclax-Based Therapy in Patients with Acute Myeloid Leukemia. Cancer Discov 10, 536–551 (2020). 10.1158/2159-8290.CD-19-0710

80 Duy, C. et al. Chemotherapy Induces Senescence-Like Resilient Cells Capable of Initiating AML Recurrence. Cancer Discov 11, 1542–1561 (2021). 10.1158/2159-8290.CD-20-1375

81 Ngo, S. et al. Acute myeloid leukemia maturation lineage influences residual disease and relapse following differentiation therapy. Nat Commun 12, 6546 (2021). 10.1038/s41467-021-26849-w

82 Boitano, A. E. et al. Aryl hydrocarbon receptor antagonists promote the expansion of human hematopoietic stem cells. Science 329, 1345–1348 (2010). 10.1126/science.1191536

83 Pabst, C. et al. Identification of small molecules that support human leukemia stem cell activity ex vivo. Nat Methods 11, 436–442 (2014). 10.1038/nmeth.2847

84 Cornet, E. et al. Involvement of a common progenitor cell in core binding factor acute myeloid leukaemia associated with mastocytosis. Leuk Res 36, 1330–1333 (2012). 10.1016/j.leukres.2012.07.001

85 Pullarkat, S. T. et al. Characterization of bone marrow mast cells in acute myeloid leukemia with t(8;21) (q22;q22); RUNX1-RUNX1T1. Leuk Res 37, 1572–1575 (2013). 10.1016/j.leukres.2013.08.010

86 Cook, J. A. et al. Fusion-harboring mast cells can explain molecular positivity in flow cytometric MRD-negative core-binding factor AML. Blood 144, 581–585 (2024). 10.1182/blood.2024024264

87 Soboli, A. et al. Presence of Leukemia-Related Basophils and Mast Cells with Atypical Immunophenotype during Induction Should Not be Interpreted As Measurable Residual Disease in Children with Acute Myeloid Leukemia with RUNX1::RUNX1T1. Blood 144, 2954 (2024). 10.1182/blood-2024-199996

88 Iwasaki, H. et al. The order of expression of transcription factors directs hierarchical specification of hematopoietic lineages. Genes Dev 20, 3010–3021 (2006). 10.1101/gad.1493506

89 Katsumura, K. R., Liu, P., Kim, J. A., Mehta, C. & Bresnick, E. H. Pathogenic GATA2 genetic variants utilize an obligate enhancer mechanism to distort a multilineage differentiation program. Proc Natl Acad Sci U S A 121, e2317147121 (2024). 10.1073/pnas.2317147121

90 Zheng, S., Papalexi, E., Butler, A., Stephenson, W. & Satija, R. Molecular transitions in early progenitors during human cord blood hematopoiesis. Mol Syst Biol 14, e8041 (2018). 10.15252/msb.20178041

