## Supplementary figures & legends, supplementary table legends for "RUNX1::RUNX1T1 Depletion Eliminates Stemness and Induces Bidirectional Differentiation of Acute Myeloid Leukemia"

#### Supplementary figure legends

**Supplementary Figure 1**. **LDV-LNPs have improved uptake and siRNA delivery efficacy in t(8;21) positive primary AML cells**. (A) Gating strategy for the LNP uptake experiments. (B) LDV-LNP uptake in t(8;21) primary AML bone marrow aspirate from patient A (left) and normal karyotype AML bone marrow aspirate (right). (C) Column graphs showing *RUNX1::RUNX1T1* fusion transcript levels in individual patient samples after sequential LNP dosing (indicated with the black arrows). (D) RUNX1::RUNX1T1 fusion protein of siRNA LNP-treated patient cells detected by western blotting 3 days after an LNP dose (patient B) or 3 days after 3 LNP doses with 3 days between sequential doses (patient D).

**Supplementary Figure 2. RUNX1::RUNX1T1 depletion leads to global chromatin changes in a t(8;21) AML PDX.** (A–B) *RUNX1::RUNX1T1* fusion transcript (A) and protein (B) levels of siRNA LNP-treated t(8;21) AML PDX cells detected by qPCR (A) or Western blotting (B). (A) Mean + ranges are displayed. n = 3 technical replicates. (C) Heatmap showing Pearson correlation of log2, counts-per-million (CPM) normalised counts between replicates and treatments, distal sites only. (D) Heatmaps depicting the chromatin accessibility in siMM and siRE-treated PDX samples, with replicates shown separately. Distal (>1.5kb from TSS) ATAC-seq sites were ranked by the fold change between siRE and siMM. The bar on the left indicates the significantly (adj.p<0.05, fold change >2) gained (red bar) and lost (blue bar) peaks. (E–G) UCSC Genome Browser screenshot showing *CTSG* (E)*, RNASE2* (F) and *VSIR* (G) loci. Upper four tracks show DNase1-seq and RUNX1::RUNX1T1 ChIP-seq [48], lower four tracks show patient C PDX ATAC-seq and RNA-seq. siRE, *RUNX1::RUNX1T1* siRNA; siMM, mismatch control.

**Supplementary Figure 3. RUNX1::RUNX1T1 depletion in PDX AML cells leads to global transcriptome changes.** (A) PCA plot of the bulk RNAseq data of the PDX AML samples treated with siRE (red) or MM (blue). N=3 biological replicates. (B–C) GSEA plots of siRNA-treated PDX samples for gene sets of down- (upper panel) and upregulated (bottom panel) RUNX1::RUNX1T1 transcriptional targets in HSCs (B, data from Tonks et al. [67]); and for gene sets of differentially (log2FC < -1 or log2FC > 1, adj. p < 0.05) down- (top panel) or upregulated (bottom panel) genes in SKNO-1 cell line upon *RUNX1::RUNX1T1* depletion with siRE (C, data from Issa et al. [21]). (D) Volcano plot of all identifiable genes in a differential expression analysis (DESeq2) of siRNA-treated PDX samples. Adjusted p-value cutoff = 0.001. Log2FC cutoff = 1. siRE, *RUNX1::RUNX1T1* siRNA; siMM, mismatch control.

**Supplementary Figure 4. *RUNX1::RUNX1T1* silencing drives differentiation of primary AML cells without affecting normal cell populations.** (A) Experimental scheme showing the workflow for the scRNAseq analysis. (B–D) UMAP plots of the scRNAseq data of co-cultures of MSC with primary t(8;21) AML cells (patients A, B) and t(8;21) AML PDX (patient C), colored by the SNP-based donor inference (B), myeloid/MSC marker gene expression levels (C): *LYZ* (upper panel), *NT5E* (middle panel), *THY1* (bottom panel), or inferred cell cycle stage (D). (E) Volcano plot showing differentially expressed genes in scRNAseq data of normal cells of the hematopoietic lineage in the 2 donors. A log2 fold change of 1 and an adjusted p-value of 0.05 were the thresholds. Wilcoxon rank with Bonferroni correction was used to determine the p-value. (F) GSEA plots for the scRNAseq of the primary t(8;21) AML cells (patients A, B) ant t(8;21) AML PDX (patient C), treated with LDV-LNP-siRE *(RUNX1::RUNX1T1* knockdown), for the gene sets of differentially (log2FC < -1 or log2FC > 1, adj.p < 0.05) expressed genes in the bulk RNAseq of the t(8;21) AML PDX (patient C) upon *RUNX1::RUNX1T1* depletion with siRE (data from the present work, Figure 3 and Supplementary Figure 3). (G) Box and violin plot of confidence scores for annotation of normal bone marrow cells from AML patient B. Wilcoxon rank sum test was used to determine the p-value. (H) PCA plot of the integrated scRNAseq dataset of the 3 patients colored by experimental condition (siRE, red and siMM, blue). siRE, *RUNX1::RUNX1T1* siRNA; siMM, mismatch control.

**Supplementary Figure 5. Depletion of RUNX1::RUNX1T1 enhances bidirectional granulocytic and eosinophilic differentiation.** (A) Box and violin plots showing cell type prediction confidence score in scRNAseq data of the 3 patients with and without RUNX1::RUNX1T1 depletion. (B) UMAP plots of integrated patient scRNAseq data, siRE *(RUNX1::RUNX1T1* knockdown) condition only, colored by module scores of gene sets from the Human Cell Atlas bone marrow gene set collection [55]: CD34-positive granulocytes (left) and CD34-positive eosinophil, basophil and mast cell progenitor (right). (C–D) STRING-generated gene networks of clusters formed by genes upregulated upon *RUNX1::RUNX1T1* silencing and higher expressed in supercluster I reflecting ribosomal components (C) and a chaperone cluster (D). (E) UMAP plot of integrated patient scRNAseq data colored by normalized expression of *CEBPE* (left), *IL5RA* (middle), *GATA2* (right). (F–G) tSNE plots of multi-color flow cytometry results of primary t(8;21) AML bone marrow aspirates, colored by patient (F) or CD125 scaled signal (G). siRE, *RUNX1::RUNX1T1* siRNA; siMM, mismatch control.

**Supplementary Figure 6. RUNX1::RUNX1T1 depletion compromises leukemic self-renewal.** (A) GSEA plots of scRNAseq data, using the HSC multipotent progenitor gene set from the cord blood gene set collection [90]. (B) tSNE plots of multi-color flow cytometry results of primary t(8;21) AML cells with or without *RUNX1::RUNX1T1* knockdown colored by scaled signal of CD34. Highlighted with blue line are cell groups enriched in the siMM condition; red, siRE condition. (C–D) tSNE plots of multi-color flow cytometry results of primary t(8;21) AML bone marrow aspirates, colored by patient (C) or scaled signals of CD34 and CD38 (D). Highlighted with blue line are cell groups enriched in the siMM condition; red, siRE condition (D). (E) UMAP plot of the integrated scRNAseq dataset of the 3 patients, colored by inferred cell cycle stage. siRE, *RUNX1::RUNX1T1* siRNA; siMM, mismatch control.

#### Supplementary tables

**Supplementary Table 1**. **Characteristics of the primary AML samples used for the study.**

**Supplementary Table 2**. **Sequences of qPCR primers used for the study.**

**Supplementary Table 3. Sequences of TotalSeq-B Hashtag Oligos used for the study.**

**Supplementary Table 4. Flow cytometry panel for cells of patient C (PDX).**

**Supplementary Table 5. Flow cytometry panel for cells of patients A, B, D.**

**Supplementary Table 6. Results of differential gene expression analysis of bulk RNAseq data of patient C (PDX), siRE *(RUNX1::RUNX1T1* knockdown) vs siMM (mismatch control).** Positive log2fold changes indicate an upregulation in the siRE condition; negative, in siMM.

**Supplementary Table 7. Results of differential gene expression analysis of scRNAseq data of patients A, B, C (PDX), by patient and all 3 patients integrated, siRE *(RUNX1::RUNX1T1* knockdown) vs siMM (mismatch control).** Positive log2fold changes indicate an upregulation in the siRE condition; negative, in siMM.

**Supplementary Table 8. Results of differential gene expression analysis of scRNAseq data of patients A, B, C (PDX), all patients integrated, supercluster I vs supercluster II, both experimental conditions (siRE and siMM) and siRE condition only.** Positive log2fold changes indicate an upregulation in supercluster I; negative, in supercluster II.

Figure S1

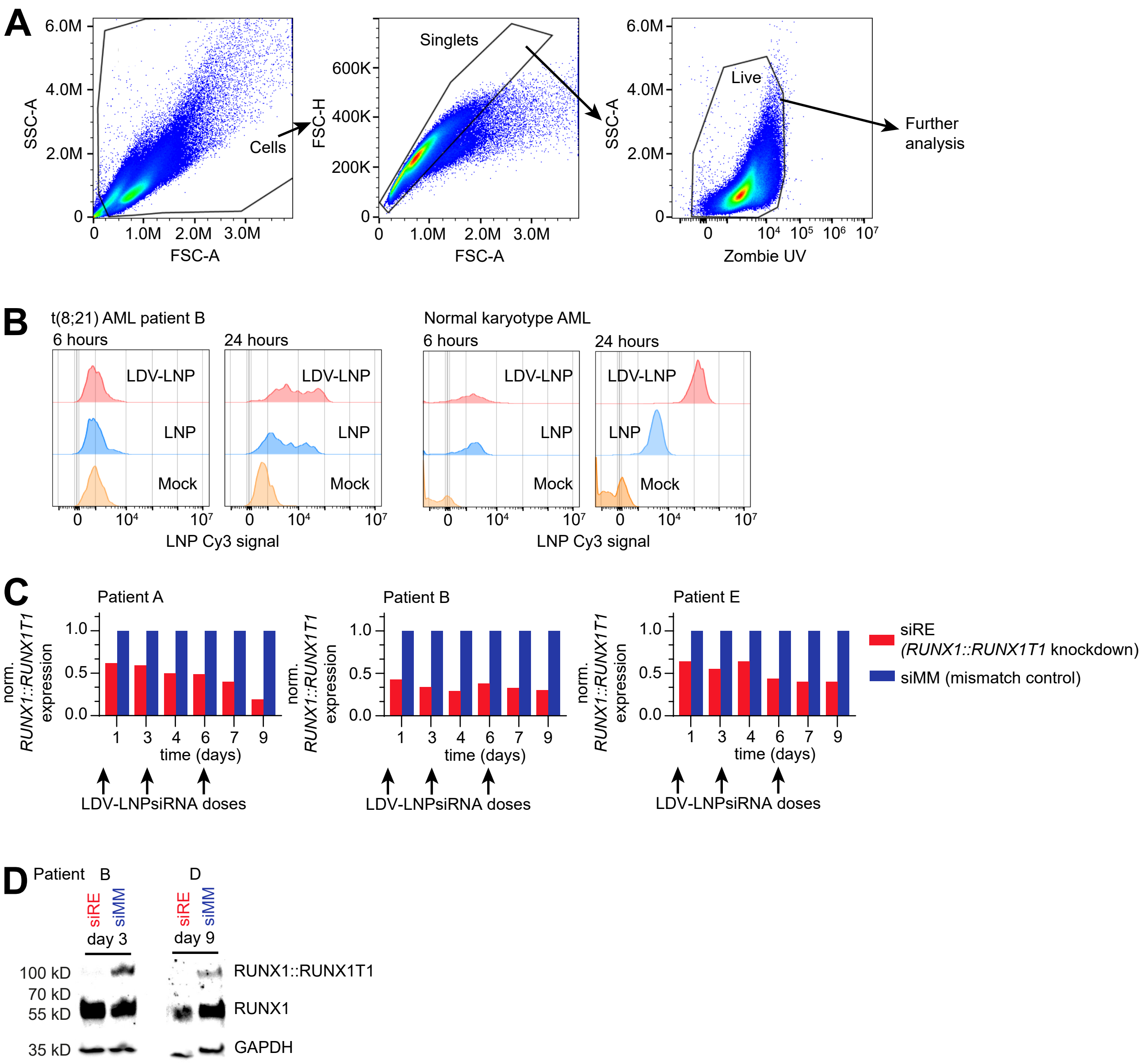
A
SSC-A
FSC-A
Cells
FSC-H
Singlets
Live
Zombie UV
Further analysis
B
t(8;21) AML patient B
Normal karyotype AML
6 hours
24 hours
LDV-LNP
LNP
Mock
LNP Cy3 signal
C
Patient A
Patient B
Patient E
norm. RUNX1::RUNX1T1 expression
time (days)
LDV-LNPsiRNA doses
siRE (RUNX1::RUNX1T1 knockdown)
siMM (mismatch control)
D
Patient B
D
siRE
siMM
day 3
day 9
100 kD
70 kD
55 kD
35 kD
RUNX1::RUNX1T1
RUNX1
GAPDH

Figure S2

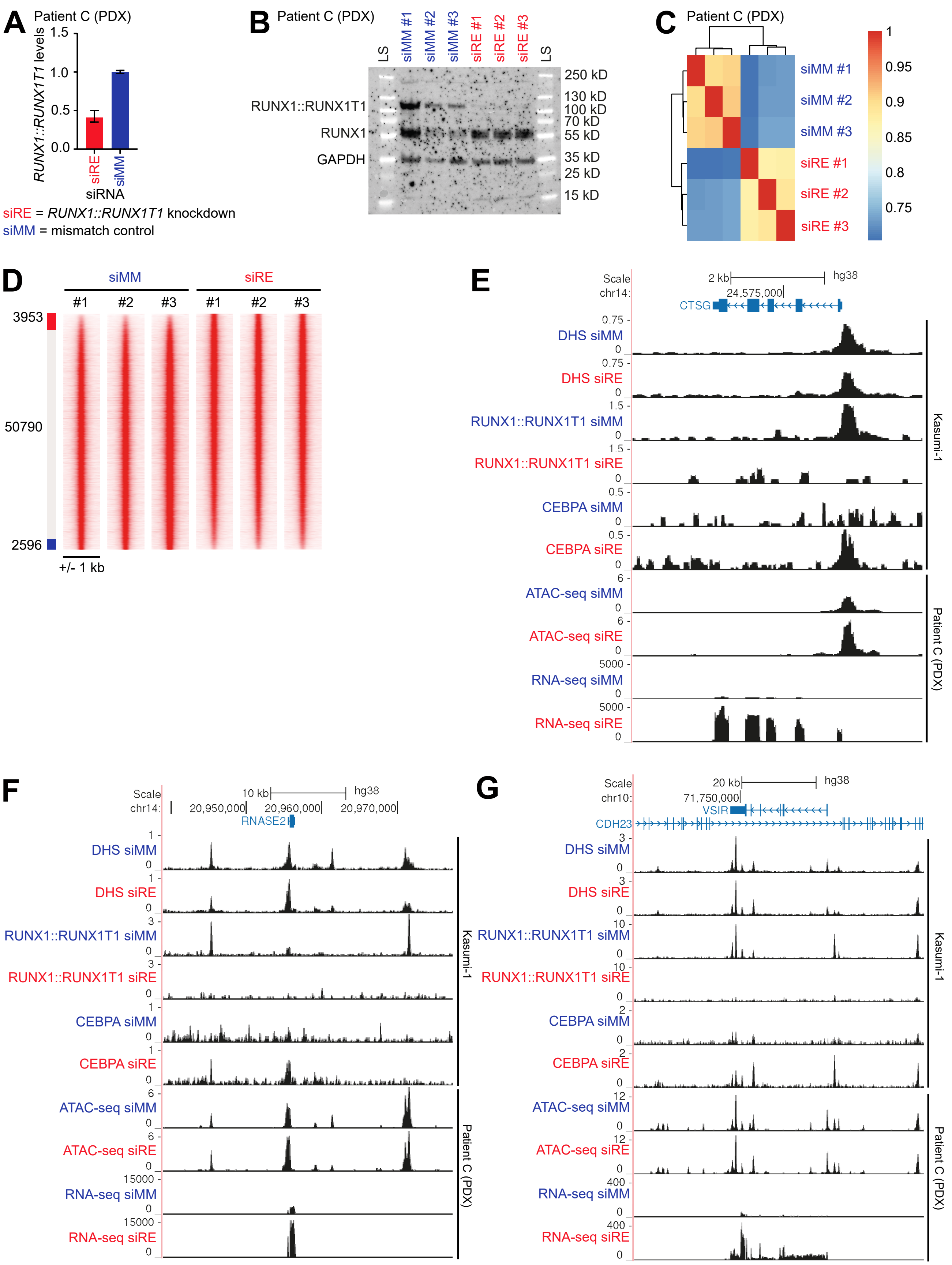
A
Patient C (PDX)
RUNX1::RUNX1T1 levels
1.5
1.0
0.5
0.0
siRE
siMM
siRNA
siRE = RUNX1::RUNX1T1 knockdown
siMM = mismatch control
B
Patient C (PDX)
LS
siMM #1
siMM #2
siMM #3
siRE #1
siRE #2
siRE #3
LS
RUNX1::RUNX1T1
RUNX1
GAPDH
250 kD
130 kD
100 kD
70 kD
55 kD
35 kD
25 kD
15 kD
C
Patient C (PDX)
siMM #1
siMM #2
siMM #3
siRE #1
siRE #2
siRE #3
1
0.95
0.9
0.85
0.8
0.75
D
siMM
siRE
#1
#2
#3
#1
#2
#3
3953
50790
2596
+/- 1 kb
E
Scale
chr14:
2 kb
hg38
24,575,000
CTSG
DHS siMM
DHS siRE
RUNX1::RUNX1T1 siMM
RUNX1::RUNX1T1 siRE
CEBPA siMM
CEBPA siRE
ATAC-seq siMM
ATAC-seq siRE
RNA-seq siMM
RNA-seq siRE
Kasumi-1
Patient C (PDX)
F
Scale
chr14:
10 kb
hg38
20,950,000
20,960,000
20,970,000
RNASE2
DHS siMM
DHS siRE
RUNX1::RUNX1T1 siMM
RUNX1::RUNX1T1 siRE
CEBPA siMM
CEBPA siRE
ATAC-seq siMM
ATAC-seq siRE
RNA-seq siMM
RNA-seq siRE
Kasumi-1
Patient C (PDX)
G
Scale
chr10:
20 kb
hg38
71,750,000
VSIR
CDH23
DHS siMM
DHS siRE
RUNX1::RUNX1T1 siMM
RUNX1::RUNX1T1 siRE
CEBPA siMM
CEBPA siRE
ATAC-seq siMM
ATAC-seq siRE
RNA-seq siMM
RNA-seq siRE
Kasumi-1
Patient C (PDX)

Figure S3

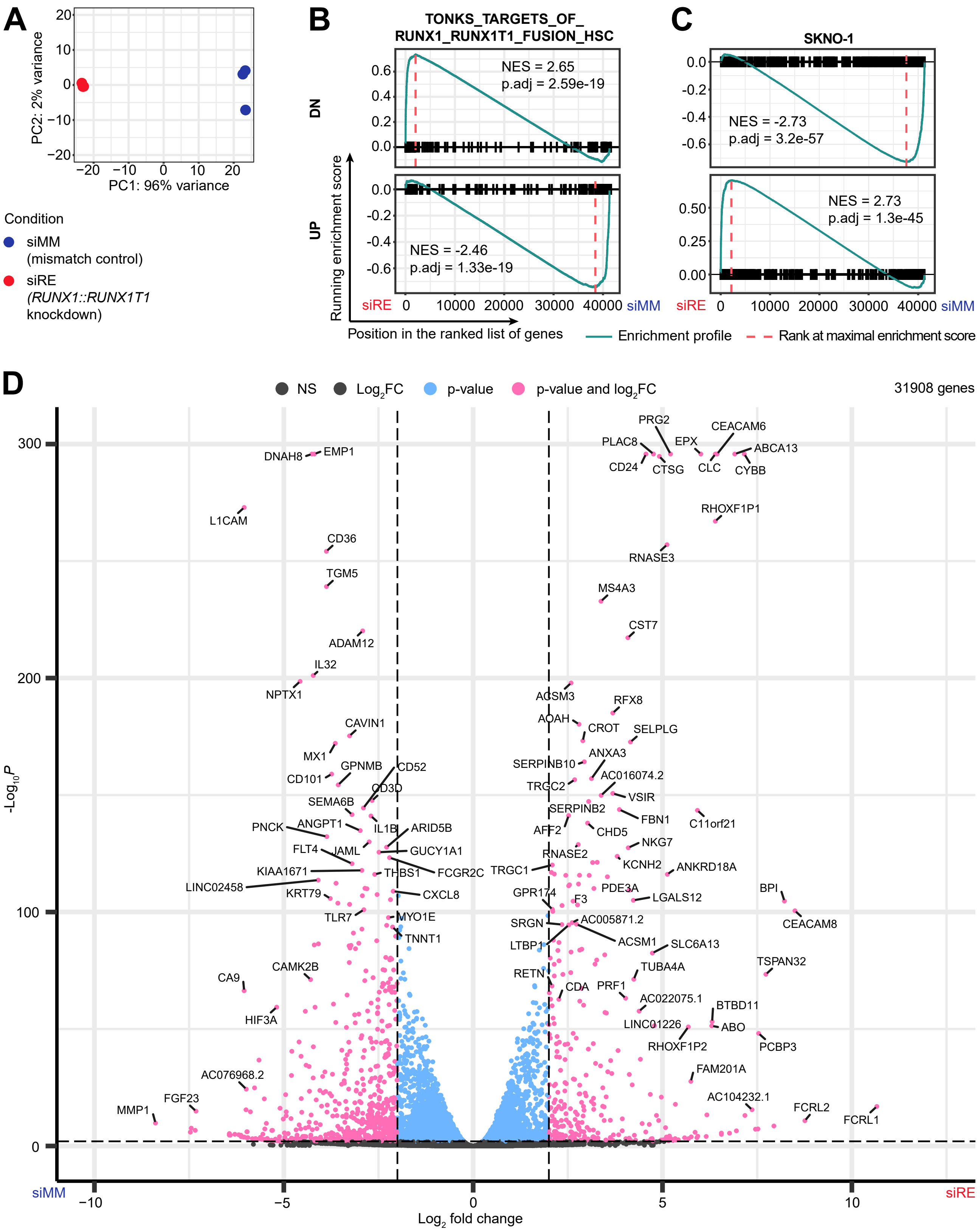

A
PC2: 2% variance
PC1: 96% variance
Condition
siMM (mismatch control)
siRE (RUNX1::RUNX1T1 knockdown)
B
TONKS_TARGETS_OF_RUNX1_RUNX1T1_FUSION_HSC
DN
NES = 2.65
p.adj = 2.59e-19
UP
NES = -2.46
p.adj = 1.33e-19
Running enrichment score
Position in the ranked list of genes
C
SKNO-1
NES = -2.73
p.adj = 3.2e-57
NES = 2.73
p.adj = 1.3e-45
siRE
siMM
Enrichment profile
Rank at maximal enrichment score
D
NS
Log2FC
p-value
p-value and log2FC
31908 genes
-Log10P
Log2 fold change
siMM
siRE

Figure S4

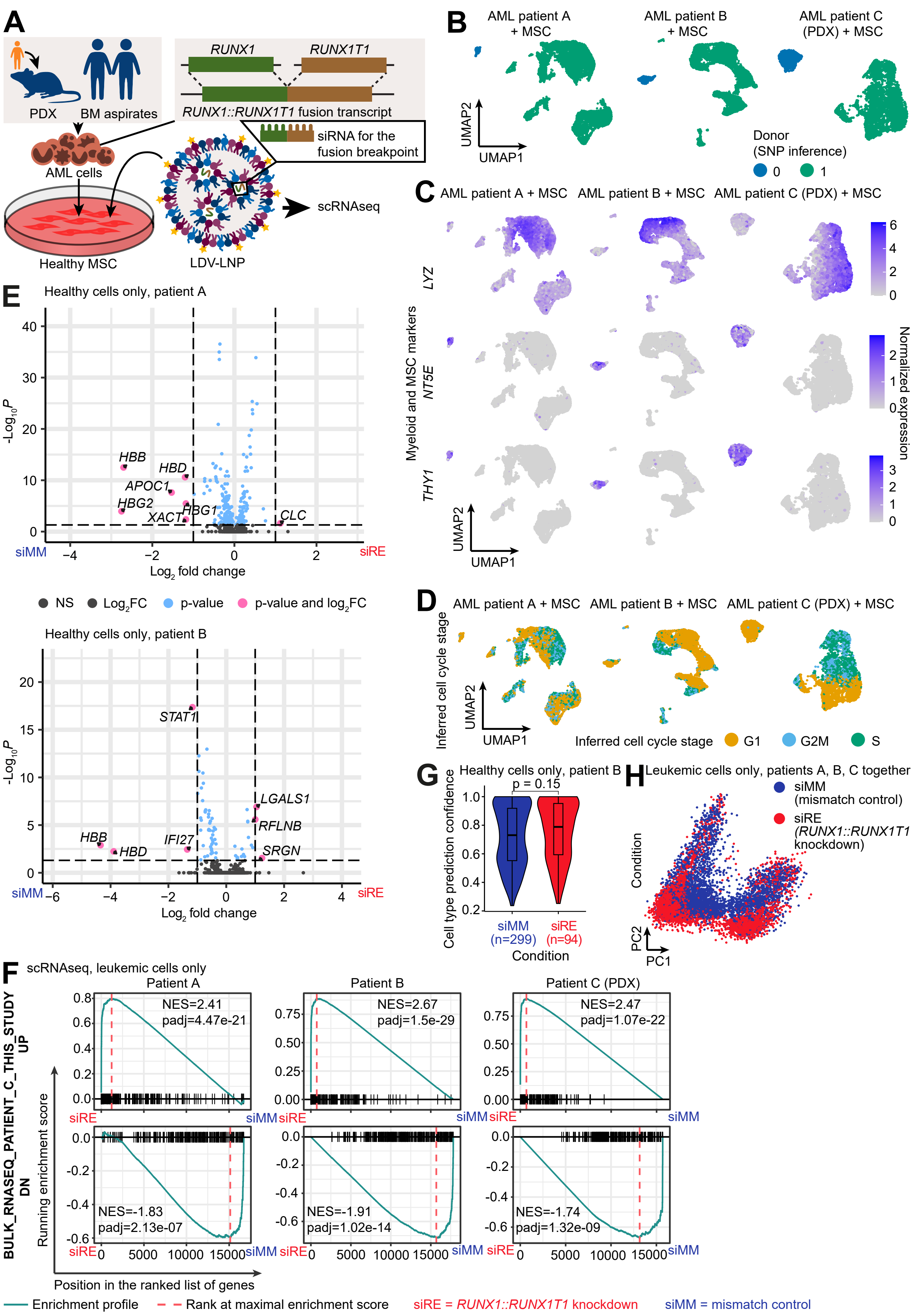
A
PDX
BM aspirates
RUNX1
RUNX1T1
RUNX1::RUNX1T1 fusion transcript
siRNA for the fusion breakpoint
AML cells
Healthy MSC
LDV-LNP
scRNAseq
B
AML patient A + MSC
AML patient B + MSC
AML patient C (PDX) + MSC
UMAP2
UMAP1
Donor (SNP inference)
0
1
C
AML patient A + MSC
AML patient B + MSC
AML patient C (PDX) + MSC
Myeloid and MSC markers
LYZ
NT5E
THY1
Normalized expression
UMAP2
UMAP1
D
AML patient A + MSC
AML patient B + MSC
AML patient C (PDX) + MSC
Inferred cell cycle stage
UMAP2
UMAP1
Inferred cell cycle stage G1 G2M S
E
Healthy cells only, patient A
-Log10 P
HBB
HBD
APOC1
HBG2
XACT
HBG1
CLC
siMM
siRE
Log2 fold change
NS
Log2FC
p-value
p-value and log2FC
Healthy cells only, patient B
STAT1
LGALS1
RFLNB
SRGN
HBB
HBD
IFI27
G
Healthy cells only, patient B
p = 0.15
Cell type prediction confidence
siMM (n=299)
siRE (n=94)
Condition
H
Leukemic cells only, patients A, B, C together
siMM (mismatch control)
siRE (RUNX1::RUNX1T1 knockdown)
Condition
PC2
PC1
F
scRNAseq, leukemic cells only
Patient A
Patient B
Patient C (PDX)
BULK_RNASEQ_PATIENT_C_THIS_STUDY
UP
DN
NES=2.41
padj=4.47e-21
NES=2.67
padj=1.5e-29
NES=2.47
padj=1.07e-22
NES=-1.83
padj=2.13e-07
NES=-1.91
padj=1.02e-14
NES=-1.74
padj=1.32e-09
Running enrichment score
Position in the ranked list of genes
siRE
siMM
Enrichment profile
Rank at maximal enrichment score
siRE = RUNX1::RUNX1T1 knockdown
siMM = mismatch control

### Figure S5

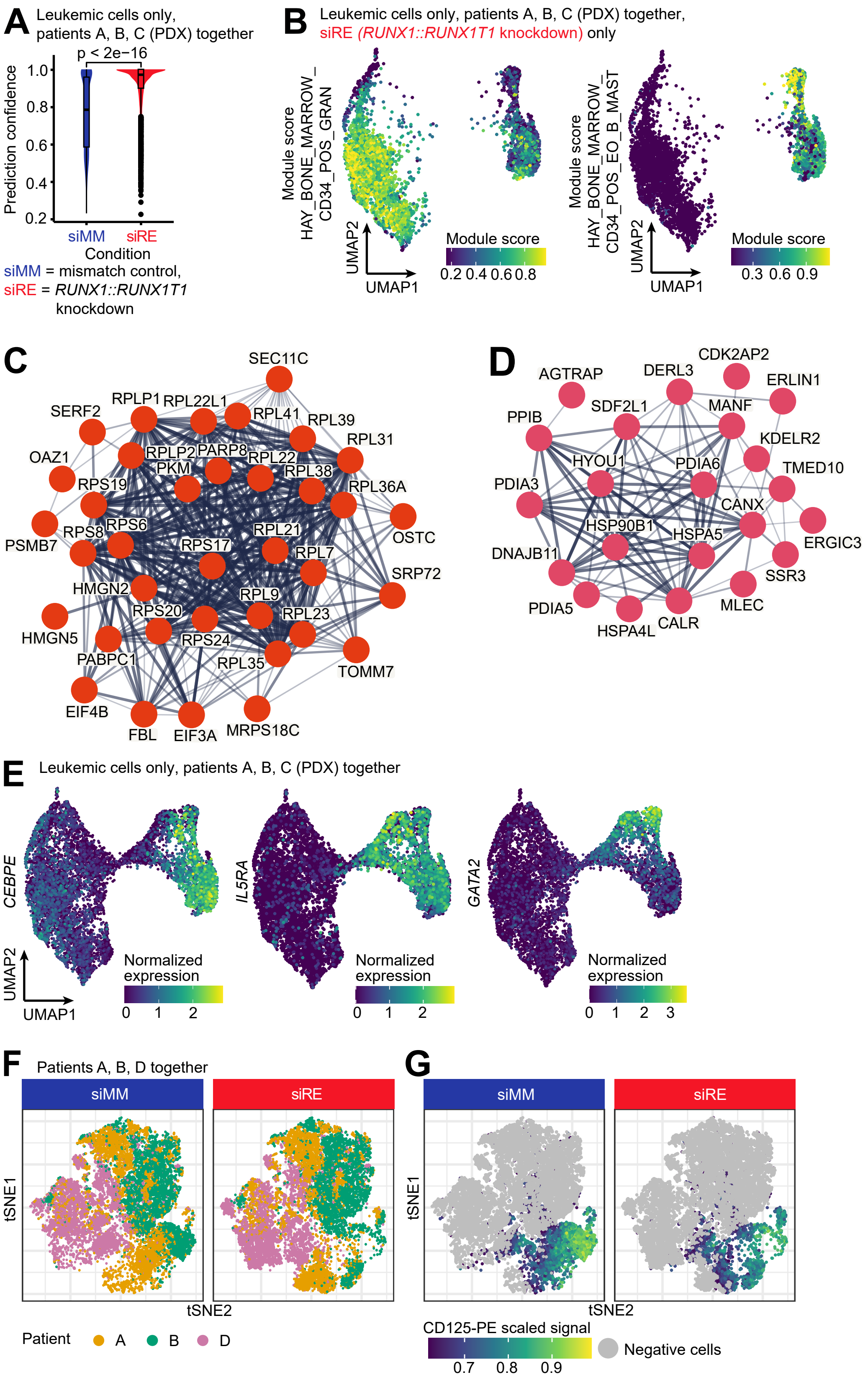
A
Leukemic cells only, patients A, B, C (PDX) together
p < 2e−16
Prediction confidence
1.0
0.8
0.6
0.4
0.2
siMM
siRE
Condition
siMM = mismatch control,
siRE = RUNX1::RUNX1T1 knockdown
B
Leukemic cells only, patients A, B, C (PDX) together,
siRE (RUNX1::RUNX1T1 knockdown) only
Module score HAY_BONE_MARROW_ CD34_POS_GRAN
Module score HAY_BONE_MARROW_ CD34_POS_EO_B_MAST
UMAP2
UMAP1
Module score
0.2 0.4 0.6 0.8
0.3 0.6 0.9
C
SEC11C
RPLP1
RPL22L1
RPL41
RPL39
SERF2
RPL31
OAZ1
RPLP2
PARP8
RPL22
PKM
RPL38
RPS19
RPL36A
PSMB7
RPS8
RPS6
OSTC
RPS17
RPL21
RPL7
SRP72
HMGN2
RPL9
RPS20
RPL23
HMGN5
RPS24
PABPC1
RPL35
TOMM7
EIF4B
FBL
EIF3A
MRPS18C
D
AGTRAP
DERL3
CDK2AP2
ERLIN1
SDF2L1
MANF
PPIB
KDELR2
HYOU1
PDIA6
TMED10
PDIA3
CANX
HSP90B1
HSPA5
ERGIC3
DNAJB11
SSR3
MLEC
PDIA5
HSPA4L
CALR
E
Leukemic cells only, patients A, B, C (PDX) together
CEBPE
IL5RA
GATA2
UMAP2
UMAP1
Normalized expression
0 1 2
0 1 2
0 1 2 3
F
Patients A, B, D together
siMM
siRE
tSNE1
tSNE2
Patient
A
B
D
G
siMM
siRE
tSNE1
tSNE2
CD125-PE scaled signal
0.7 0.8 0.9
Negative cells

### Figure S6

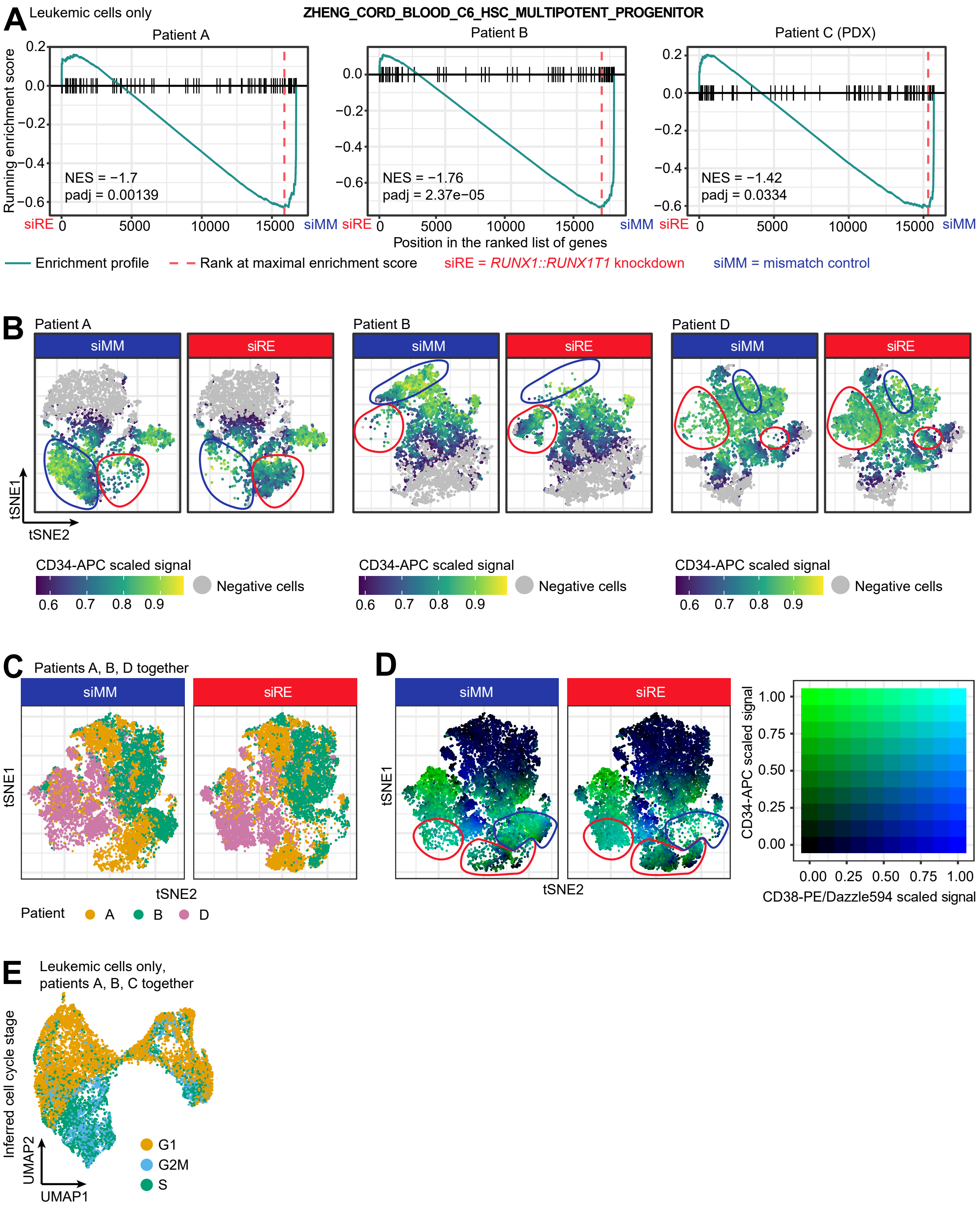

A
Leukemic cells only
ZHENG_CORD_BLOOD_C6_HSC_MULTIPOTENT_PROGENITOR
Patient A
Patient B
Patient C (PDX)
Running enrichment score
NES = −1.7
padj = 0.00139
NES = −1.76
padj = 2.37e−05
NES = −1.42
padj = 0.0334
siRE
siMM
Position in the ranked list of genes
Enrichment profile
Rank at maximal enrichment score
siRE = RUNX1::RUNX1T1 knockdown
siMM = mismatch control
B
Patient A
Patient B
Patient D
siMM
siRE
tSNE1
tSNE2
CD34-APC scaled signal
Negative cells
0.6 0.7 0.8 0.9
C
Patients A, B, D together
Patient A B D
D
CD34-APC scaled signal
CD38-PE/Dazzle594 scaled signal
0.00 0.25 0.50 0.75 1.00
E
Leukemic cells only,
patients A, B, C together
Inferred cell cycle stage
UMAP1
UMAP2
G1
G2M
S
